# *Caenorhabditis Elegans* Junctophilin has Tissue-Specific Functions and Regulates Neurotransmission with Extended-Synaptotagmin

**DOI:** 10.1101/2020.11.20.392142

**Authors:** Christopher A. Piggott, Zilu Wu, Stephen Nurrish, Suhong Xu, Joshua M. Kaplan, Andrew D. Chisholm, Yishi Jin

## Abstract

The junctophilin family of proteins tether together plasma membrane (PM) and endoplasmic reticulum (ER) membranes, and couple PM- and ER-localized calcium channels. Understanding *in vivo* functions of junctophilins is of great interest for dissecting the physiological roles of ER-PM contact sites. Here, we show that the sole *C. elegans* junctophilin JPH-1 localizes to discrete membrane contact sites in neurons and muscles and has important tissue-specific functions. *jph-1* null mutants display slow growth and development due to weaker contraction of pharyngeal muscles, leading to reduced feeding. In the body wall muscle, JPH-1 co-localizes with the PM-localized EGL-19 voltage-gated calcium channel and ER-localized UNC-68/RyR calcium channel, and is required for animal movement. We also find an unexpected cell non-autonomous effect of *jph-1* in axon regrowth after injury. In neurons, JPH-1 co-localizes with the membrane contact site protein Extended-SYnaptoTagmin 2 (ESYT-2) and modulates neurotransmission. Interestingly, *jph-1* and *esyt-2* null mutants display mutual suppression in their response to aldicarb, suggesting that JPH-1 and ESYT-1 have antagonistic roles in neuromuscular synaptic transmission. Our genetic double mutant analysis also reveals that *jph-1* functions in overlapping pathways with two PM-localized voltage-gated calcium channels, *egl-19* and *unc-2,* and *unc-68/RyR* for animal health and development. Finally, we show that *unc-68/RyR* is required for JPH-1 localization to ER-PM puncta. Our data demonstrate important roles for junctophilin in cellular physiology, and also provide insights into how junctophilin functions together with other calcium channels *in vivo*.

## Introduction

Membrane contact sites (MCSs) are regions of close contact, generally within 10 to 30 nm between organelles or between an organelle and the plasma membrane (PM). MCSs were first described between the endoplasmic reticulum (ER) and PM in muscle cells by electron microscopy over 60 years ago (Porter and Palade, 1957). MCSs have now been found for most organelles in many organisms (Lang et al., 2015; Valm et al., 2017). MCSs are maintained by protein tethers that bind to opposing membranes simultaneously and hold them in close proximity. Different types of MCSs are organized by distinct protein tethers, many of which are conserved from yeast to mammals (Phillips and Voeltz, 2016). Recent studies have begun to uncover their functions. For example, oxysterol-binding proteins (OSBPs) facilitate exchange of PM-localized phosphatidylinositol 4-phosphate (PI4P) for ER-localized cholesterol (Mesmin et al., 2013), and binding of ER-localized calcium sensor Stim1 to PM-localized calcium channel Orai1 triggers the entry of extracellular calcium to the ER to replenish calcium stores (Hirve et al., 2018). Genetic analysis suggests many MCS tethering proteins act redundantly. For example, studies of ER-PM contact sites in yeast showed that full separation of the ER from the PM is only achieved when six genes encoding MCS proteins are deleted (Manford et al., 2012). Similarly, in *C. elegans,* enlarged lysosomes and endosomes were observed only when knocking out all four *obr* genes encoding OSBP homologs (Kobuna et al., 2010). It is thus necessary to identify new experimental models or paradigms to tease apart the functions of individual MCS proteins and to dissect their interaction network *in vivo.*

The junctophilin (JPH) family of proteins were first identified based on their localization to muscle ER-PM contact sites in a screen using monoclonal antibodies raised against ER vesicles enriched for ER-PM junctions(Takeshima et al., 2000). Junctophilins are characterized by a N-terminal domain consisting of eight <u>m</u>embrane <u>o</u>ccupation and <u>r</u>ecognition <u>n</u>exus (MORN) motifs, which bind to the PM, and a C-terminal transmembrane domain, which anchors the protein to the ER. Mammals have four junctophilins (JPH1 through 4) that are differentially expressed in excitable cells. JPH1 and JPH2 are expressed in skeletal and cardiac muscle (Nishi et al., 2000; Takeshima et al., 2000) and the smooth muscle surrounding arteries (Pritchard et al., 2019; Saeki et al., 2019). JPH3 and JPH4 are broadly expressed in neurons of the brain and many parts of the nervous system (Nishi et al., 2003, 2000; Takeshima et al., 2000). Studies of genetic knockout mice have provided some evidence for their functions. Cardiomyocytes from JPH2 knockout mice have fewer ER-PM contacts, and skeletal muscle from JPH1 knockout mice have abnormal ER morphology and fewer ER-PM contacts (Ito et al., 2001; Takeshima et al., 2000). In addition to tethering together ER and PM membranes, junctophilins bind to ER-and PM-localized calcium channels and facilitate their co-localization at ER-PM contact sites in mouse cardiomyocytes, skeletal muscle, and cultured hippocampal neurons (Nakada et al., 2018; Sahu et al., 2019; Van Oort et al., 2011). Junctophilin-mediated ER-PM coupling is reported to promote efficient excitation-contraction in heart and skeletal muscle (Ito et al., 2001; Nakada et al., 2018; Takeshima et al., 2000; Van Oort et al., 2011) and regulate action potential frequency in neurons (Kakizawa et al., 2007; Moriguchi et al., 2006; Sahu et al., 2019). Unlike mammals, invertebrates have a single junctophilin (Garbino et al., 2009). In *D. melanogaster,* the sole junctophilin was shown to have roles in muscle contraction and neural development (Calpena et al., 2018).

*C. elegans* has a single junctophilin gene named *jph-1* (Garbino et al., 2009; Yoshida et al., 2001). Here we show that JPH-1 protein localizes to punctate structures in muscles and neurons. In muscles, JPH-1 puncta co-localize with the ER-localized UNC-68/RyR calcium channel and PM-localized EGL-19/Cav1 calcium channel. In neurons JPH-1 puncta co-localize with the ER-PM contact site protein extended-synaptotagmin 2 (ESYT-2). Through characterization of *jph-1* null mutants and tissue-specific rescue experiments, we defined tissue-specific roles of *jph-1.* In the pharynx muscle, *jph-1* is required for the pumping that drives animal feeding and contributes to animal growth. In the body wall muscle, *jph-1* is required for animal movement. We observed a cell non-autonomous effect of *jph-1* in axon regeneration after injury. Additionally, *jph-1* modulates synaptic transmission, and can balance the effects of *esyt-2.* Genetic double mutant analyses reveal differential interactions between *jph-1* and the ER-localized *unc-68/RyR* calcium channel and two PM-localized voltage-gated calcium channels (VGCCs) for animal development and health. Lastly, we show that precise localization of JPH-1 in both neurons and muscles depends on *unc-68.* These data support critical roles of junctophilin in cellular function and animal development.

## Materials and methods

### *C. elegans* genetics

Wild-type *C. elegans* is the N2 Bristol variant (Brenner, 1974). Strains were maintained under standard conditions on Nematode Growth Media (NGM) plates seeded with *E. coli* OP50 bacteria. New strains were constructed using standard procedures, based on a combination of visual identification of phenotypes, such as uncoordinated (Unc) movement, and genotyping for specific alleles. Strains and primers for genotyping are shown in the reagents table.

### Molecular biology and transgenesis

We cloned *jph-1* cDNAs from wild-type N2 mRNAs, first using primers YJ12558 5’-GACGTAGGTGTGTCAGCAG-3’ and YJ12559 5’-CCTGAGGAGAAGTGTGTCTG-3’ in the 5’UTR and 3’UTR of *jph-1,* followed by a second round of amplification using primers YJ12560 5’-ATGAATGGAGGCAGATTTGAC-3’ and YJ12561 5’-CTACGAAGAAGACTTCTTCTTCTTC-3’ targeting the start and stop codons. We obtained two amplified products, which were cloned into pCR8 vectors. Sanger sequencing analysis of these clones revealed a 2.2 kb cDNA encoding JPH-1 isoform A, and a 2.4 kb cDNA encoding JPH-1 isoform B. The coding region of JPH-1B was then amplified using primers YJ12560 5’-ATGAATGGAGGCAGATTTGAC-3’ and YJ12562 5’-CTAATATGTGAGGGTGTGTACCG-3’ and cloned into a pCR8 vector. The 4.5 kb *jph-1* promoter was amplified from wild-type genomic DNA using the primers YJ12563 5’-TGTTCTGCCATTACCAGCCCG-3’ and YJ12564 5’-TTCCCATTTGCCGTACTGCTG-3’. All expression constructs were generated either by Gateway recombination (Invitrogen / Thermo Fisher Scientific), Gibson assembly (New England Biolabs), or restriction enzyme digest and ligation. All expression clones were sequenced to ensure sequence fidelity.

We generated transgenic lines by microinjection, as described (Mello et al., 1991). Plasmids, fosmids, co-injection markers, and injection concentrations are listed in the reagents table.

Single-copy insertion transgenes with *ju* designation were generated on Chromosome IV at cxTi10882, following a previously published protocol (Andrusiak et al., 2019). Briefly, we injected N2 hermaphrodites with four plasmids, one containing GFP-cDNA flanked by homology arms and expressing a hygromycin resistance gene (HygR), pCZGY2750 expressing Cas9 and an sgRNA targeting cxTi10882, and pCFJ90 Pmyo-2-mCherry (Addgene plasmid 19327) and pCFJ104 Pmyo-3-mCherry (Addgene plasmid 19328)(Frøkjær-Jensen et al., 2008) as co-injection markers. F1 animals from injected P0 parents were treated with hygromycin (Hyg). Among the survivors, we looked for the absence of co-injections markers to identify animals with genomic insertion, which was further verified by PCR genotyping using primers YJ10503, YJ10504, and YJ10686 (wild type 562 bp, insertion 744 bp). Single-copy insertion transgene *nuTi144* was generated by using a modified Mos1 transposon, following a previously published protocol (Frøkjær-Jensen et al., 2014).

### CRISPR-Cas9 gene editing

We generated the *jph-1(ju1683)* and *jph-1(ju1684)* deletion alleles using two CRISPR RNAs (crRNAs): 5’-CCGTCCGGTAACACCTATCA-3’ and 5’-ACGACGTTGACCAGCAAGAC-3’ (Integrated DNA Technologies) targeting *jph-1* exon 1 and exon 9, respectively. The crRNAs were injected into wild-type hermaphrodites with purified Cas9 (MacroLabs, University of California, Berkeley), trans-activating crRNA (tracrRNA) and *dpy-10* crRNA, as described (Paix et al., 2015). We selected small and slow-growing Unc animals resembling *jph-1(ok2823)* mutants, as we were unable to isolate animals based on Dpy or Rol phenotypes, possibly because the *jph-1* crRNA was more efficient than the *dpy-10* crRNA. We identified *ju1683* and *ju1684* as deletions in *jph-1* by PCR genotyping with flanking primers YJ12565 5’-GACGACGGCGGAACCTATG-3’ and YJ12566 5’-TCAGGTACGTTCTAGTCGGT-3’.

GFP11 knock-in alleles *unc-68(nu664)* and *egl-19(nu674)* were generated by injecting wild-type hermaphrodites with 75 ng/μl pDD162 expressing Cas9, 36 ng/μl pRB1017-derived guide RNA, and 75ng/ul of a PCR product of 7 copies of GFP11 flanked by 1 kb of wild-type sequence 5’ and 3’ of the cut site. Guide RNAs were selected using the CRISPR guide RNA selection tool (http://genome.sfu.ca/crispr/). A gRNA targeting *unc-58* (pGW28) 36 ng/μl and repair oligo (AF-JA-76) were also injected as a co-conversion marker (El Mouridi et al., 2017).

### Animal growth assessment

Adult hermaphrodite animals were placed on seeded NGM plates and allowed to lay eggs for two hours, after which they were removed. The plates were kept at 20°C and observed daily to determine the time it took the offspring to reach the fourth larval (L4) stage.

### Brood size assay

L4 hermaphrodite animals were individually placed on seeded NGM plates and moved to new plates daily. Two days after a parent animal was placed on a plate, the number of hatched offspring were counted. This was continued until parent animals laid no more eggs or died. The number of hatched offspring produced per parent animal was totaled to calculate brood size.

### Fluorescence microscopy

Animals were immobilized in a drop of M9 solution with or without 30 mM muscimol or 10 mM levamisole on a 4% agar pad or 10% agarose pad. Most confocal fluorescence images were collected using a Zeiss LSM800 confocal microscope with Z-stacks taken at 0.5 or 1 μm intervals between planes for most images, with the exception of 0.21 μm intervals for GFP::JPH-1A in *unc-68(0)* (**Figure 7A,B**). *Pjph-1-GFP* in the head and tail (**Supplemental Figure 2A**) were imaged using a Zeiss LSM710 confocal microscope, and GFP::JPH-1A in the PLM neuron (**Supplemental Figure 4B**) was imaged using an Andor spinning disk confocal unit (CSU-W1) with a Leica DMi8 microscope. All confocal fluorescence images were taken at 63x magnification. Maximum intensity projections were prepared using Fiji (ImageJ).

Images of GFP-labeled touch neurons [P*mec-7*-GFP(*muls32*)j in wild-type and *jph-1(0)* animals were taken on a Zeiss Axio Imager A2 compound scope at 10x magnification under identical settings.

### Brightfield microscopy

Images depicting gross body morphology (**Figure 1C**) were taken by immobilizing animals in a drop of M9 solution on a 10% agarose pad and imaging on a Leica DMi8 microscope under brightfield settings at 10x magnification with an Andor iXon Ultra camera. Images depicting animal movement crawling on NGM petri plates (**Figure 4A**) were taken on a Zeiss M2 stereodissecting microscope with a Nikon DS-Qi1Mc camera.

**Figure 1.**
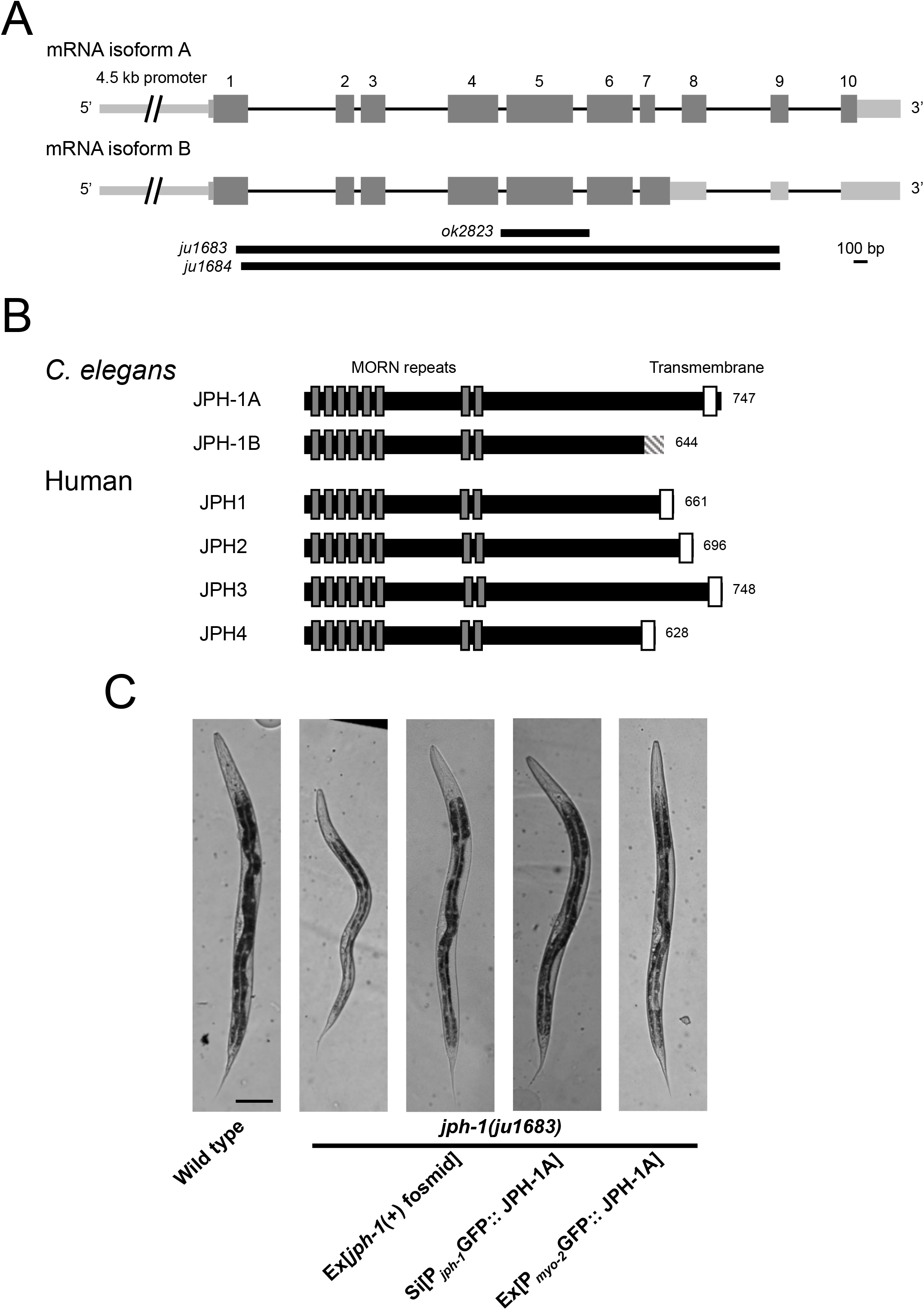
*jph-1* expresses two isoforms that differ at their C-termini and is required for animal development. **A**) Illustration of *jph-1* spliced isoforms and deletion alleles. Exons are dark grey boxes, introns are black lines, and UTRs are light grey boxes. *ok2823* is a 637 bp deletion, *ju1683* is a 3891 bp deletion, and *ju1684* is a 3858 bp deletion with a 13 bp insertion. **B**) Illustration of *C. elegans* JPH-1 proteins predicted from isolated cDNA sequences in comparison to human JPH proteins. Dark grey boxes indicate membrane occupation and recognition nexus (MORN) repeats and white boxes indicate transmembrane domains. The striped box at the C-terminus of JPH-1B indicates the 35 amino residues predicted from the cDNA. These 35 amino acids are not predicted to form a transmembrane region or low-complexity domain using Pfam (El-Gebali et al., 2019), the TMHMM Server v 2.0 (http://www.cbs.dtu.dk/services/TMHMM/), or SMART (Letunic and Bork, 2018). A BLASTp search of these 35 amino acids against all published *Caenorhabditis* genomes (Caenorhabditis.org) also revealed no significant hits with a low e-value threshold of 1.0. Gene accession numbers are: JPH-1A (NP_492193.2), Human JPH1 (NP_001304759.1), JPH2 (NP_065166.2), JPH3 (NP_065706.2), and JPH4 (NP_001139500.1). Pairwise sequence alignments were performed between *C. elegans* JPH-1A and human JPH1, JPH2, JPH3, and JPH4 using MUSCLE (Madeira et al., 2019) and the Percent Identity Matrix was viewed to find percent identity. To determine conservation between MORN repeats, we concatenated all eight 14 amino acid MORN repeats into one sequence for each protein and then performed pairwise sequence alignments using MUSCLE. Sequence identity ranges from 69% to 77% when comparing only MORN sequences in *C. elegans* JPH-1A and human JPH1 through 4. **C**) Bright field images of L4 stage animals of genotypes indicated. Compared to wild type animals *jph-1(ju1683)* animals are small, thin, and pale, all of which was rescued by transgenic expression of a fosmid containing genomic *jph-1 (juEx3390),* JPH-1A expressed under the *jph-1* promoter *[Pjph-1-GFP::JPH-1A(juSi387)],* or JPH-1A expressed in the pharyngeal muscle *[Pmyo-2-GFP::JPH-1A(juEx8041)].* Scale bar 100 μm.

### Pharyngeal pumping assays

To count pumping rate, day-1 adult animals on seeded plates were observed through dissection stereomicroscopes. We counted the number of grinder movements in 20 seconds twice per animal and took the average. Counting was done while animals were on the OP50 bacterial lawn to prevent variations in pumping rate caused by food availability.

To measure pumping strength, we adapted a published protocol that used serotonin to stimulate pumping in immobilized animals (Trojanowski and Fang-Yen, 2015). We prepared 8% agarose pads with 8mM serotonin (H7752, Sigma Aldrich), placed animals in an M9 drop on the pad, and immediately placed a cover slip on top. We began imaging when animals started pumping (about 0-10 minutes after animals were placed in the M9 drop). Imaging was performed on a Leica DMi8 microscope at 40x magnification. 20 second videos were taken at 100 ms/frame for a total of 200 frames per animal. Videos were then analyzed in Fiji (ImageJ). The distance from the grinder to an arbitrary point on the pharyngeal lumen was measured in the frame immediately preceding pump initiation (**Figure 3B, left image**). The distance from the grinder to the same point was measured in the frame when the grinder had moved to its fullest extent (**Figure 3B, right image**). The difference between these two measurements is the distance moved by the grinder in one pump. We took the average of the first five pumps in each video, although in three instances wild-type animals only pumped three or four times during the video. The distance moved by the grinder was divided by the length of the pharynx (**Figure 3B, left image**) to normalize to animal size.

### Thrashing assay

Individual L4 animals were placed in 1 μl drops of M9 on a glass dissection plate. We counted the number of thrashes performed by the animal in one minute. We considered a single thrash to be one sufficiently large movement of the animal’s head or tail back and forth, with the head or tail not necessarily crossing the centre of mass.

### Aldicarb and levamisole assays

To test aldicarb sensitivity, 15 day-1 adult animals were transferred to fresh plates containing 0.5 mM or 1 mM aldicarb. Animals were scored for paralysis every 30 minutes by gently touching the animal with a platinum wire. For levamisole sensitivity, 15 day-1 adult animals were transferred to fresh plates containing 1 mM levamisole. Animals were scored for paralysis every 15 minutes by gently touching the animal with a platinum wire. Final sample size for each assay was 13-15 animals due to some animals crawling off the plate. Drug sensitivity was quantified from three independent experiments.

### Laser axotomy of PLM axons

We cut PLM axons and quantified the length of regrown axons as previously described (Wu et al., 2007). Briefly, GFP-labeled PLM axons [P*mec-7*-GFP(*muls32*)] of L4 animals were cut 40 μm anterior to the cell body by a femtosecond laser on a spinning-disk confocal microscope. Animals were recovered onto seeded NGM plates and the regrown axon was imaged 24 hours later on a Zeiss LSM510 or LSM800 confocal microscope.

### Statistical analysis

We used Prism (GraphPad Software) for all statistical analysis except for Fisher’s exact test, for which we used the online tool QuickCalcs (Graphpad Software). To compare regrowth between experiments with different control means, we normalized each experimental data point by dividing it by its control means. Statistical tests and sample sizes are indicated in Figures or Figure legends.

### Strains used in this manuscript

**Strains used in this manuscript:**
| Strain | Genotype |
| --- | --- |
| N2 | wild type |
| CZ27358 | *jph-1(ju1683) I* |
| CZ27360 | *jph-1(ju1684) I* |
| CZ28073 | *jph-1(ju1683) I; juEx3390[jph-1 fosmid WRM0622aB02]* |
| CZ28260 | *jph-1(ju1683) I; juEx3392[jph-1 fosmid WRM0623aF07]* |
| CZ27162 | *juEx8014[Pjph-1-GFP]* |
| CZ27161 | *juEx8013[Pjph-1-GFP]* |
| CZ28024 | *jph-1(ju1683) I; juEx7999[Pjph-1-GFP::JPH-1A]* |
| CZ27777 | *jph-1(ju1683) I; juSi387[Pjph-1-GFP::JPH-1A] IV* |
| CZ27603 | *jph-1(ju1683) I; juEx8037[Pjph-1-GFP::JPH-1B]* |
| CZ27606 | *jph-1(ju1683) I; juEx8038[Pjph-1-GFP::JPH-1B]* |
| CZ27364 | *juSi387[Pjph-1-GFP::JPH-1A] IV* |
| CZ28264 | *nuTi144 [Pmyo-3 GFP 1-10 G418] I; unc-68(nu664) V; juEx8103[Pjph-1-mKate2::JPH-1A]* |
| CZ28263 | *nuTi144 [Pmyo-3 GFP 1-10 G418] I; egl-19(nu674) IV; juEx8103[Pjph-1-mKate2::JPH-1A]* |
| CZ27404 | *juSi387[Pjph-1-GFP::JPH-1A] IV; juls540[Pmec-4-mKate2::ESYT-2] X* |
| CZ27569 | *juEx8038[Pjph-1-GFP::JPH-1B]* |
| CZ24806 | *eat-4(ky5)* |
| CZ27782 | *jph-1(ju1683) I; juEx8041[Pmyo-2-GFP::JPH-1A]* |
| CZ28120 | *jph-1(ju1683) I; juEx8022[Pmyo-3-GFP::JPH-1A]* |
| CZ28121 | *jph-1(ju1683) I; juEx8023[Pmyo-3-GFP::JPH-1A]* |
| CZ28051 | *jph-1(ju1683) I; juSi389(Prab-3-GFP::JPH-1A) IV* |
| CZ13170 | *jph-1(ok2823) I* |
| CZ13990 | *jph-1(ok2823) I; juEx3390[jph-1 fosmid WRM0622aB02]* |
| CZ27483 | *jph-1(ok2823) I; juEx3392[jph-1 fosmid WRM0623aF07]* |
| CZ10969 | *muls32(Pmec-7-GFP) II* |
| CZ26391 | *jph-1(ok2823) I; muls32[Pmec-7-GFP] II* |
| CZ27359 | *jph-1(ju1683) I; muls32[Pmec-7-GFP] II* |
| CZ27778 | *jph-1(ju1683) I; muls32[Pmec-7-GFP] II; juSi387[Pjph-1-GFP::JPH-1A] IV* |
| CZ27783 | *jph-1(ju1683) I; muls32[Pmec-7-GFP] II; juEx8041[Pmyo-2-GFP::JPH-1A]* |
| CZ27910 | *jph-1(ju1683) I; muls32[Pmec-7-GFP] II; juSi388[Pmec-4-GFP::JPH-1A] IV* |
| OD2984 | *ltSi953[Pmec-18-Degron-SL2-mKate2; cb-unc-119(+)] II; unc-119(ed3) III* |
| CZ27842 | *jph-1(ju1683) I; ltSi953[Pmec-18-Degron-SL2-mKate2; cb-unc-119(+)] II; juSi387[Pjph-1-GFP::JPH-1A] IV; unc-119(ed3)? III* |
| CZ27361 | *jph-1(ju1684) I; muls32[Pmec-7-GFP] II* |
| CZ27536 | *juEx8035[Pjph-1-GFP::JPH-1(ok2823)]* |
| CZ28262 | *nuls321[Punc-17-mCherry]; juEx7999[Pjph-1-GFP::JPH-1A]* |
| CZ27875 | *juSi388[Pmec-4-GFP::JPH-1A] IV* |
| CZ26322 | *esyt-2(ju1409) III* |
| CZ28238 | *esyt-2(ju1409) III; juEx7581 [esyt-2 gDNA]* |
| CZ28070 | *jph-1(ju1683) I; esyt-2(ju1409) III* |
| CZ26389 | *esyt-2(ju1408) III* |
| CZ28241 | *jph-1(ju1684) I; esyt-2(ju1408) III* |
| CZ28282 | *jph-1(ju1683) I; esyt-2(ju1409) III; juEx7581 [esyt-2 gDNA]* |
| CB540 | *unc-68(e540) V* |
| CZ28026 | *jph-1(ju1683) I; unc-68(e540)V; juEx7999[Pjph-1-GFP::JPH-1A]* |
| CZ26691 | *egl-19(ad1006lf) IV* |
| DA695 | *egl-19(ad695gf) IV* |
| CZ27905 | *jph-1(ju1683) I; egl-19(ad695gf) IV* |
| CB55 | *unc-2(e55) X* |
| CZ28054 | *jph-1(ju1683) I; unc-2(e55) X; juEx7999[Pjph-1-GFP::JPH-1A]* |
| QW47 | *unc-2(zf35gf) X* |
| CZ27903 | *jph-1(ju1683) I; unc-2(zf35gf) X* |
| CZ28025 | *unc-68(e540) V; juEx7999[Pjph-1-GFP::JPH-1A]* |
| CZ28053 | *unc-2(e55) X; juEx7999[Pjph-1-GFP::JPH-1A]* |
| CZ28055 | *esyt-2(ju1409) III; juSi387[Pjph-1-GFP::JPH-1A] IV* |

### Plasmid used in this manuscript

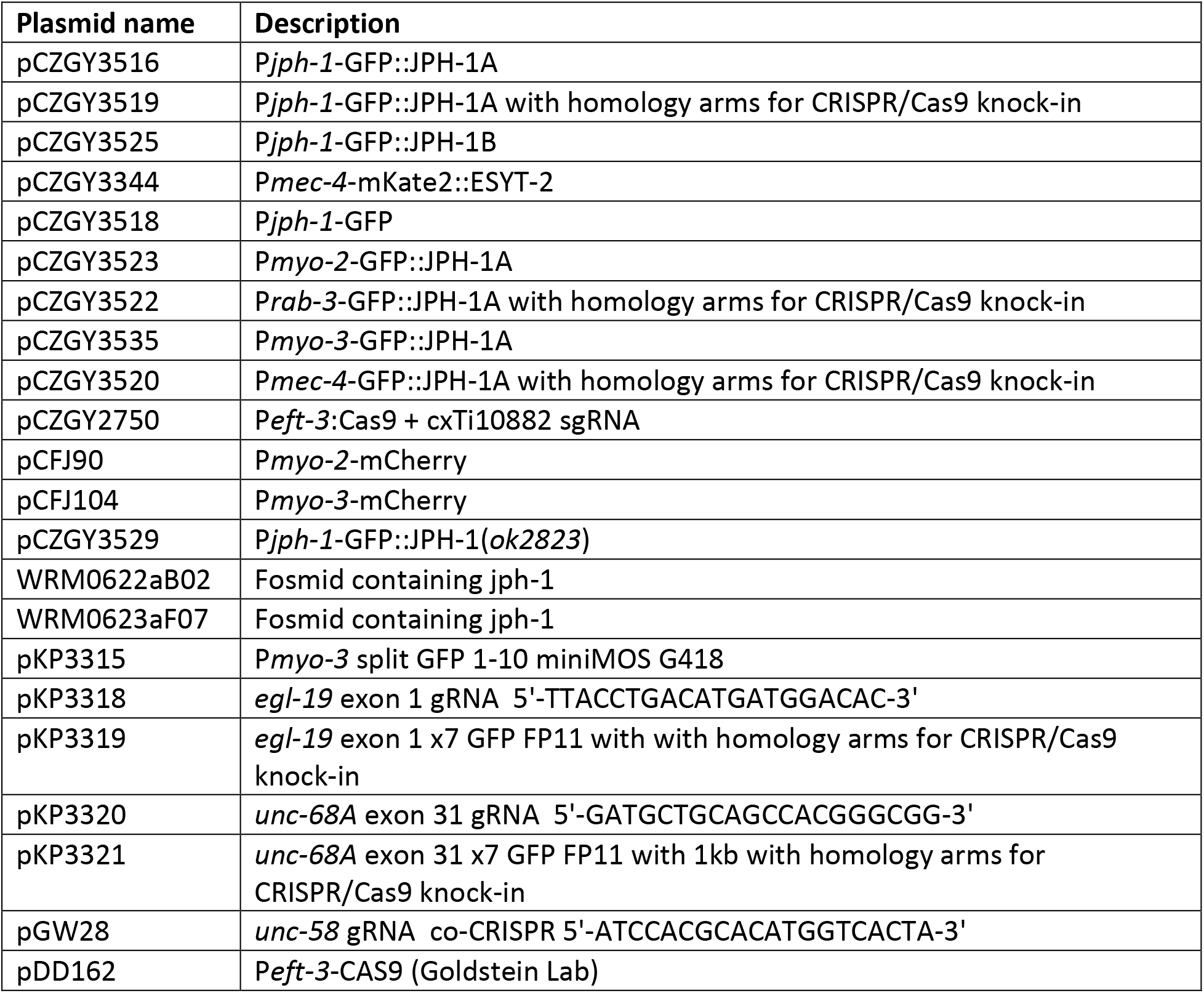

### Transgenes used in this manuscript

**Transgenes used in this manuscript:**
| Transgene name | Injected DNA | Injected conc. (ng/μL) | Coinjection marker and conc. | Source |
| --- | --- | --- | --- | --- |
| *juEx3390[jph-1 fosmid WRM0622aB02]* | fosmid WRM0622aB02 | 10 | *Pttx-3*-RFP (30 ng/μL) | This manuscript |
| *juEx3392[jph-1 fosmid WRM0623aF07]* | fosmid WRM0623aF07 | 10 | *Pttx-3*-RFP (30 ng/μL) | This manuscript |
| *juEx8014[Pjph-1-GFP]* | pCZGY3518 | 2.5 | *Punc-122*-RFP (50 ng/μL) | This manuscript |
| *juEx8013[Pjph-1-GFP]* | pCZGY3518 | 2.5 | *Punc-122*-RFP (50 ng/μL) | This manuscript |
| *juEx7999[Pjph-1-GFP::JPH-1A]* | pCZGY3516 | 2.5 | *Punc-122*-RFP (50 ng/μL) | This manuscript |
| *juSi387[Pjph-1-GFP::JPH-1A]* | pCZGY3519 | 2.5 |  | This manuscript |
| *juEx8037[Pjph-1-GFP::JPH-1B]* | pCZGY3525 | 2.5 | *Punc-122*-RFP (50 ng/μL) | This manuscript |
| *juEx8038[Pjph-1-GFP::JPH-1B]* | pCZGY3525 | 2.5 | *Punc-122*-RFP (50 ng/μL) | This manuscript |
| *juls540[Pmec-4-mKate2::ESYT-2]* |  |  |  | Kim *et al.*, 2018 |
| *juEx8041[Pmyo-2-GFP::JPH-1A]* | pCZGY3523 | 0.125 | *Punc-122*-RFP (50 ng/μL) | This manuscript |
| *juEx8022[Pmyo-3-GFP::JPH-1A]* | pCZGY3535 | 2.5 | *Punc-122*-RFP (50 ng/μL) | This manuscript |
| *juEx8023[Pmyo-3-GFP::JPH-1A]* | pCZGY3535 | 2.5 | *Punc-122*-RFP (50 ng/μL) | This manuscript |
| *juSi389[Prab-3-GFP::JPH-1A]* | pCZGY3522 | 10 |  | This manuscript |
| *juSi388[Pmec-4-GFP::JPH-1A]* | pCZGY3520 | 2.5 |  | This manuscript |
| *ItSi953[pOD2087/pSW408]* |  |  |  | Wang *et al.*, 2017 |
| *juEx8035[Pjph-1-GFP::JPH-1(ok2823)]* | pCZGY3529 | 2.5 | *Punc-122*-RFP (50 ng/μL) | This manuscript |
| *nuTi144* | pKP3315 | 10 |  | This manuscript |

### Primers used in this manuscript

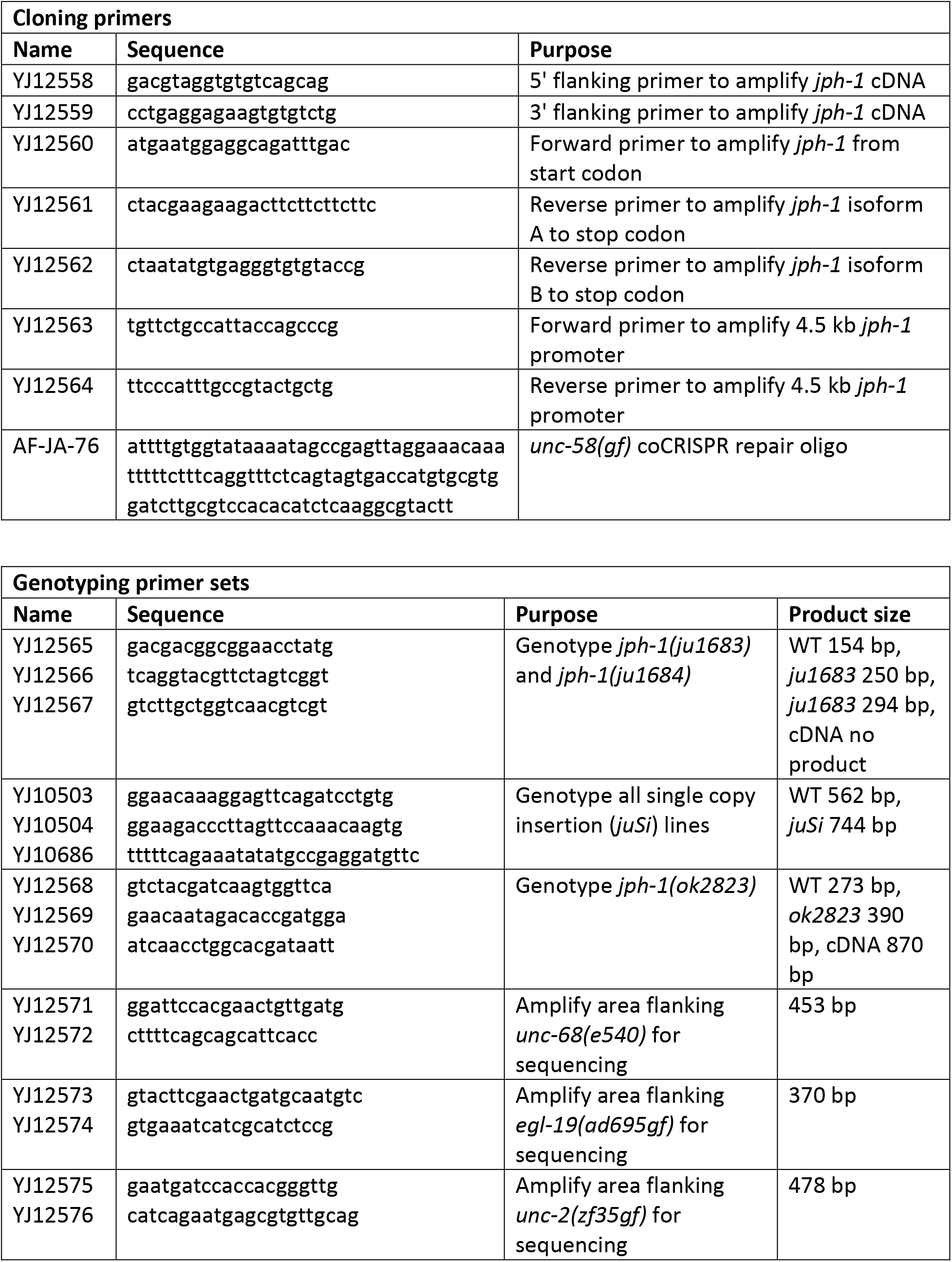
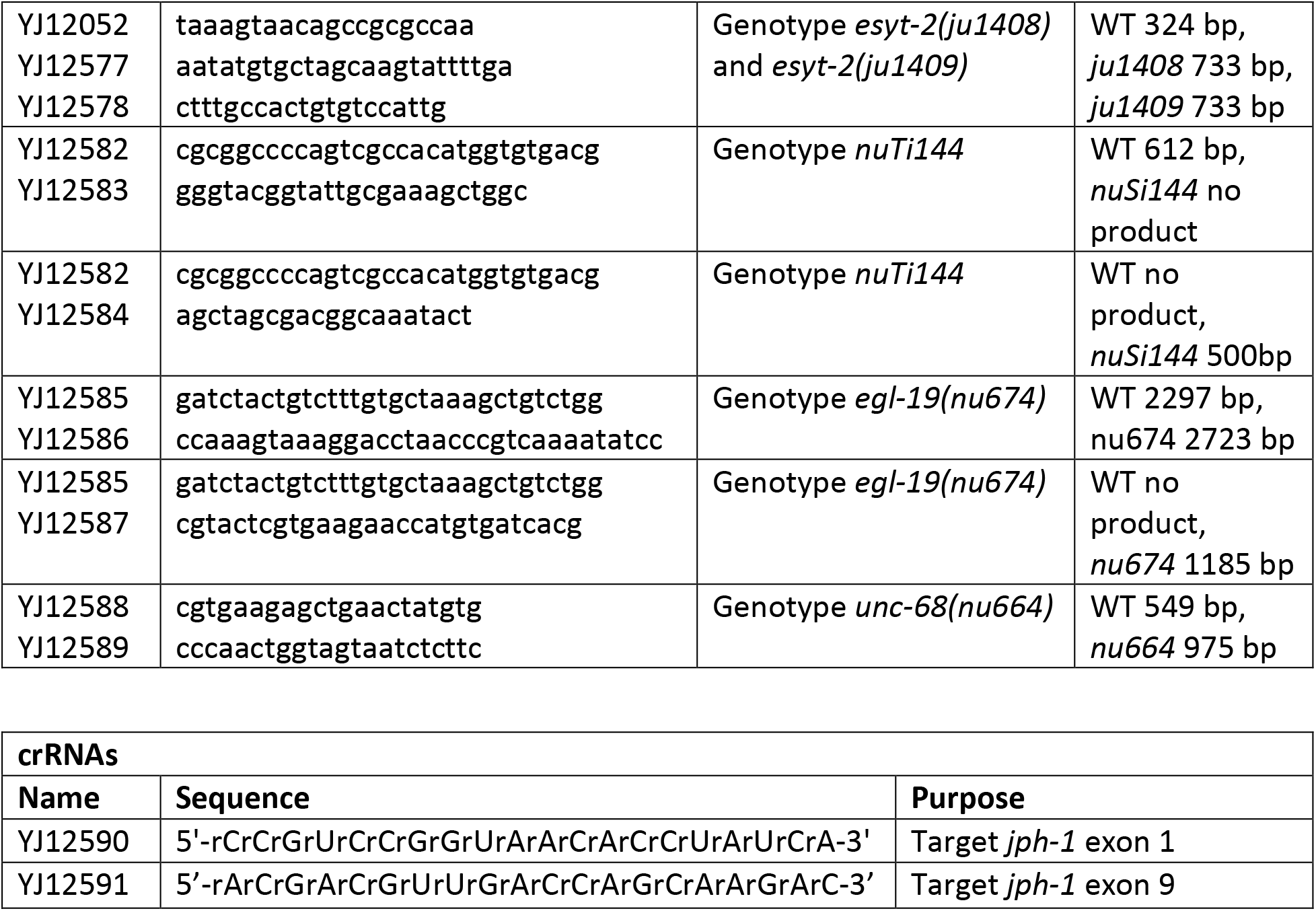

Strains and plasmids are available upon request. The authors affirm that all data necessary for confirming the conclusions of the article are present within the article, figures, and tables.

## Results

### *jph-1* expresses two isoforms

A previous study described a *jph-1* cDNA that encodes a 747 amino acid protein (Yoshida et al., 2001). In the process of obtaining *jph-1* cDNA for our own study, we obtained two cDNAs of 2.2 kb and 2.4 kb in size, amplified using primers flanking the start and stop codons (**Figure 1A**). The 2.2 kb cDNA matches the previously reported *jph-1* cDNA, which we designated isoform A (Yoshida et al., 2001). The 2.4 kb cDNA retains the intron between exon 7 and exon 8 and would encode a protein with the C-terminal 138 amino acids of the previously reported *jph-1* cDNA replaced by a different 35 amino acid sequence (**Figure 1B**). We designated this shorter protein isoform B. The C-terminal 35 amino acids do not contain a predicted transmembrane domain, nor conserved domains or low complexity regions. A BLASTp search of all published *Caenorhabditis* genomes found no significant hits for the 35 amino acid sequence, suggesting that it is not conserved. Furthermore, a BLASTn search showed that although the intron is conserved in 10 out of 26 published *Caenorhabditis* genomes (**Supplemental Figure 1A**), the translated sequences have low amino acid conservation and highly variable sequence length due to intronic stop codons (**Supplemental Figure 1B**). The absence of conserved motifs and the lack of conservation between species suggests that these 35 amino acids may not be important for the function of *jph-1.* JPH-1 isoform A shares between 39% and 42% overall sequence identity with human JPH1 through 4, with higher sequence homology in the MORN repeats, which target junctophilin to the PM (Takeshima et al., 2000).

### *jph-1* is required for normal development

To define the function of *jph-1,* we generated two null *(0)* alleles, *ju1683* and *ju1684,* using CRISPR-Cas9 editing. Both alleles delete the entire coding sequence of both *jph-1* isoforms (**Figure 1A**). These two alleles show indistinguishable phenotypes in all analyses; therefore, we generally present the quantification data for *ju1683* (**Table 1**). By gross body morphology, *jph-1(0)* mutant animals are smaller and thinner than stage-matched wild-type animals (**Figure 1C**). *jph-1(0)* mutants develop more slowly compared to wild-type animals (**Table 1**). *jph-1(0)* mutants have reduced fertility, with a brood size (52 ± 25, n = 12) about 20% of that in wildtype animals (279 ± 19, n = 10). Transgenic expression of a fosmid containing the entire *jph-1* locus rescued the developmental defects (**Table 1**). These observations indicate that *jph-1* is necessary for proper animal development.

**Table 1.** Summary of growth phenotypes of *jph-1* mutants and relevant transgenic animals

| Genotype | Transgene | Days to L4 (20°C) | Body size at L4 |
| --- | --- | --- | --- |
| wild type | none | 2 | Normal |
| *jph-1(ju1683)* | none | 3 to 4 | Small |
| *jph-1(ju1683)* | Fosmid with genomic *jph-1*<sup>1</sup> | 2 | Normal |
| *jph-1(ju1683)* | JPH-1A<sup>2</sup> | 2 | Normal |
| *jph-1(ju1683)* | JPH-1B<sup>3</sup> | 3 to 5 | Small |
| *eat-4(ky5)* | None | 2 | Normal |
| *jph-1(ju1683)* | JPH-1A in pharyngeal muscle<sup>2</sup> | 2 | Normal |
| *jph-1(ju1683)* | JPH-1A in body wall muscle<sup>2</sup> | 3 to 4 | Small |
| *jph-1(ju1683)* | JPH-1A in neurons<sup>2</sup> | 3 to 4 | Small |
| *jph-1(ok2823)* | None | 3 to 4 | Small |
| *jph-1(ok2823)* | Fosmid with genomic *jph-1*<sup>1</sup> | 2 | Normal |

<sup>1</sup> a fosmid containing the entire genomic locus of *jph-1* [WRM0622aB02(*juEx3390*) or WRM0623aF07(*juEx3392*)]
<sup>2</sup> transgenic expression of JPH-1A under its own promoter [*Pjph-1*-GFP::*JPH-1A(juSi387* or *juEx7999)*], in pharyngeal muscle [*Pmyo-2*-GFP::*JPH-1A(juEx8041)*], in body wall muscle [*Pmyo-3*-GFP::*JPH-1A(juEx8022* or *juEx8023)*], in neurons [*Prab-3*-GFP::*JPH-1A(juSi389)*]
<sup>3</sup>JPH-1B [*Pjph-1*-GFP::*JPH-1B(juEx8037* or *juEx8038)*].

### *jph-1* is expressed in muscles and neurons, and its function requires the transmembrane domain

A previously reported *jph-1* transcriptional reporter showed expression in most muscles and some neurons in the head (Yoshida et al., 2001). We made a similar transcriptional reporter using a 4.5 kb *jph-1* promoter to control GFP expression (**Supplemental Figure 2A**). We confirmed GFP expression in all hermaphrodite muscle types, including body wall, pharyngeal, vulval, uterine, stomatointestinal, anal sphincter and anal depressor muscles, with the exception of the contractile gonadal sheath (**Supplemental Figure 2A**). We also observed expression in many neurons from head to tail.

All developmental defects of *jph-1(0)* were rescued by expression of N-terminally GFP-tagged JPH-1A under the control of the 4.5 kb *jph-1* promoter (**Supplemental Figure 2B**) as either a multicopy extrachromosomal array or a single copy insertion line (**Table 1, Figure 1C**). This result indicates that GFP-tagged JPH-1A can perform the developmental functions of *jph-1* and that the 4.5 kb promoter provides sufficient tissue specificity for *jph-1* function. In contrast, transgenic expression of GFP::JPH-1B, the truncated isoform lacking the transmembrane domain, under the same promoter, did not show rescuing activity (**Supplemental Figure 2C, Table 1**). This analysis supports a conclusion that the transmembrane domain is necessary for the function of JPH-1.

### JPH-1A localizes to subcellular puncta and co-localizes with the ER-PM contact site protein ESYT-2 in neurons

We observed that the functional GFP-tagged JPH-1A showed a punctate subcellular pattern in muscles and neurons. In body wall muscle, GFP::JPH-1A localizes to rows of puncta that follow the obliquely striated pattern of the muscle (**Figure 2A,B**). In the pharyngeal muscle, JPH-1A localizes to puncta radiating from the pharyngeal lumen and lining the pharynx periphery (**Figure 2A**). We observed broad expression in neurons in the head and tail, including the bundled neuronal processes of the nerve ring, the ventral cord neurons, and touch receptor neurons (**Figure 2A-E**). In neuronal cell bodies of the head ganglia (**Figure 2C**), tail ganglia (**Figure 2D**), and ventral nerve cord (**Figure 2E**), GFP::JPH-1A shows a reticulate localization pattern and forms bright puncta near the periphery of the cell body. This localization is observed in newly hatched L1 animals and adults, suggesting that the localization is established prior to hatching and maintained into adulthood (**Supplemental Figure 2B**).

**Figure 2.**
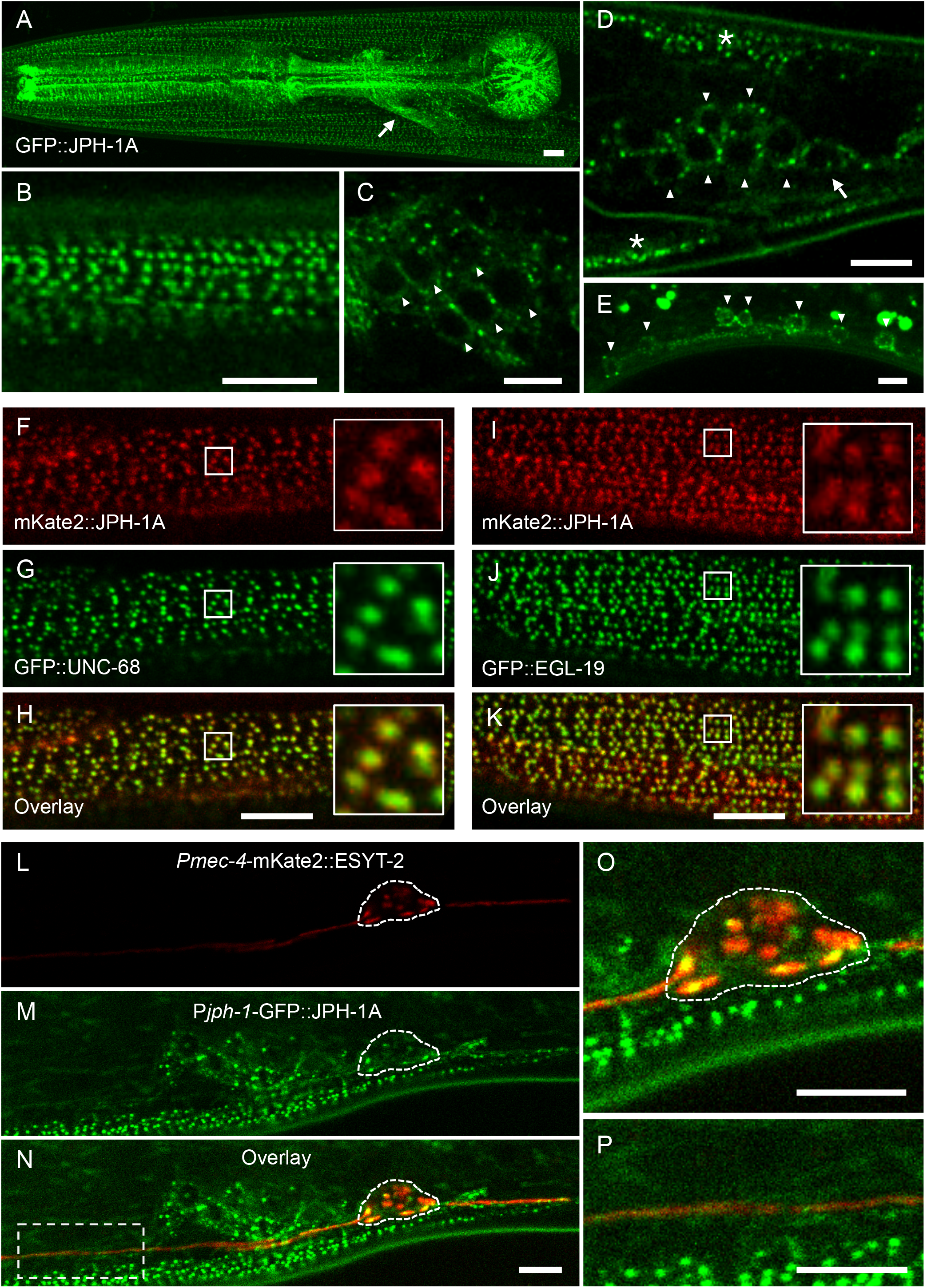
JPH-1A co-localizes with calcium channels UNC-68 and EGL-19 in muscles and MCS protein ESYT-2 in neurons. **A-D**: Confocal images of GFP::JPH-1A expressed under the *jph-1* promoter as a single copy insertion *[Pjph-1-GFP::JPH-1A(juSi387)]* in L4 stage animals. **A**) Maximum intensity projection of the head showing GFP::JPH-1A expression in body wall muscle, pharynx muscle, and neurons. Arrow indicates nerve ring. **B**) Single plane image of body wall muscle. JPH-1A localizes to rows of dots that run parallel to muscle striations. **C**) Single plane image of head ganglia neurons. JPH-1A in neuronal cell bodies is excluded from the nucleus and is concentrated in puncta. Arrowheads indicate some of the neurons expressing GFP::JPH-1A. **D**) Single plane image of tail ganglia. Arrowheads indicate neurons expressing GFP::JPH-1A. Arrow indicates PLM cell body. Asterisks mark body wall muscle. **E**) Maximum intensity projection of GFP::JPH-1A *[Pjph-1-GFP::JPH-1A(juEx7999)]* in the ventral nerve cord in an L4 stage *jph-1(ju1683)* animal. Arrowheads indicate neuronal cell bodies. Fluorescent blobs outside the cells are autofluorescent particles in the gut. **F-H**: JPH-1A co-localizes with UNC-68 in body wall muscle. Single plane confocal images of an L4 stage animal with split-GFP knock-in *unc-68 (nu664)* expressing muscle GFP1-10 [P*myo-3*-GFP1-*10(nuSi144)]* and mKate2::JPH-1A expressed under the *jph-1* promoter [P*jph-1*-mKate2::JPH-1A(*juEx8103*)]. **I-K**: JPH-1A co-localizes with EGL-19. Single plane confocal images of an L4 stage animal with split-GFP knock-in *egl-19 (nu674)* expressing muscle GFP1-10 [P*myo-3*-GFP1-10(*nuSi144*)] and mKate2::JPH-1A expressed under the *jph-1* promoter *[Pjph-1-mKate2::JPH-1A(juEx8103)].* **L-N**: JPH-1A localizes to ER-PM contact sites labeled by ESYT-2 in the cell body. Single plane confocal images of an L4 animal expressing mKate2-tagged ESYT-2 under the *mec-4* touch neuron specific promoter *[Pmec-4-mKate2::ESYT-2(juIs540)]* and GFP::JPH-1A under the *jph-1* promoter *[Pjph-1-GFP::JPH-1A(juSi387)].* PLM cell body outlined by dashed line. **O**) Close-up of panel N showing partial colocalization of JPH-1A and ESYT-2 in the PLM cell body. **P**) Close up of the box in Panel N shows that both ESYT-2 and JPH-1A are in the PLM axon. In all images, anterior is to the left, dorsal is up. Scale bars, 5 μm.

Junctophilins generally function to couple calcium channels between the ER and PM, including ER-localized ryanodine receptors (RyRs) and PM-localized L-type calcium channels (Landstrom et al., 2014). In *C. elegans, unc-68* encodes the RyR and *egl-19* encodes the Cav1 VGCC α1-subunit. We generated split-GFP knock-in lines for both *unc-68* and *egl-19* and visualized their subcellular localization by expressing muscle GFP1-10 and mKate2::JPH-1A expressed under the *jph-1* promoter. In the body wall muscle, both UNC-68 and EGL-19 localize to rows of puncta, which nearly completely overlap with JPH-1A puncta (**Figure 2F-K**). JPH-1A co-localization with both the ER-localized UNC-68 and PM-localized EGL-19 is consistent with targeting to ER-PM contact sites in muscle cells.

To determine if the neuronal puncta of JPH-1A represent MCSs, we analyzed animals coexpressing GFP::JPH-1A with a reporter line expressing mKate2::ESYT-2 in touch receptor neurons. E-Syt (extended-synaptotagmin) proteins are conserved tethering proteins at ER-PM contact sites (Giordano et al., 2013). We showed previously that *C. elegans* ESYT-2 is expressed broadly in neurons and co-localizes with an ER marker at the cell periphery (Kim et al., 2018). In the PLM soma, GFP::JPH-1A puncta co-localize with mKate2::ESYT-2 (**Figure 2L-P**), suggesting that JPH-1A clusters at ER-PM contact sites in neuronal cell bodies. We also examined GFP-tagged JPH-1B and observed a mostly diffuse localization in the muscles and neurons (**Supplemental Figure 2C**), consistent with the transmembrane domain being critical for JPH-1 subcellular localization. The lack of *jph-1(0)* rescuing activity by JPH-1B suggests that the transmembrane domain is important for its localization and function.

### *jph-1* regulates pharyngeal muscle contraction

The gross phenotypes of *jph-1(0)* mutants broadly resemble those of mutants with feeding defects in that they are small, thin, pale, and take longer to reach adulthood than wild-type animals (Avery, 1993; Avery and Horvitz, 1989). Our observation that *jph-1* is expressed in the pharynx suggests that the *jph-1(0)* phenotype may be due to defects in feeding related function. *C. elegans* eat by drawing bacteria into their mouth using pharynx pumping and crushing the bacteria with their grinder (Avery and You, 2012). We measured pumping rate by counting grinder movements and found that *jph-1(0)* mutants had a lower pumping rate than wild-type animals (**Figure 3A**). Pharyngeal muscle contraction is regulated by glutamatergic transmission. Loss of function in *eat-4,* encoding the sole glutamate transporter, causes reduced pumping rate (Lee et al., 1999). We found that *jph-1(0)* mutants had a similar pumping rate to *eat-4(ky5)* mutants (**Figure 3A**). However, since *eat-4(ky5)* animals are not as small as *jph-1(0)* mutants (**Table 1**), reduced pumping rate alone cannot account for the starved appearance of *jph-1(0)* mutants. We next quantified pumping strength by measuring the distance moved by the grinder in one pump (**Figure 3B, Materials and Methods**). *jph-1(0)* mutants had significantly weaker pumping strength than either wild type or *eat-4* mutants (**Figure 3C**). To test if reduced pharynx muscle activity was causing the starved appearance of *jph-1(0)* mutants, we expressed JPH-1A specifically in the pharynx muscle using the *myo-2* promoter. Pharyngeal muscle expression of JPH-1A restored pumping strength to wild-type levels (**Figure 3C**). Importantly, it also rescued the small body size and delayed development of *jph-1(0)* mutants (**Table 1, Figure 1C**). These observations indicate that JPH-1A is required for proper pharyngeal muscle function which ultimately impacts gross organismal development.

**Figure 3.**
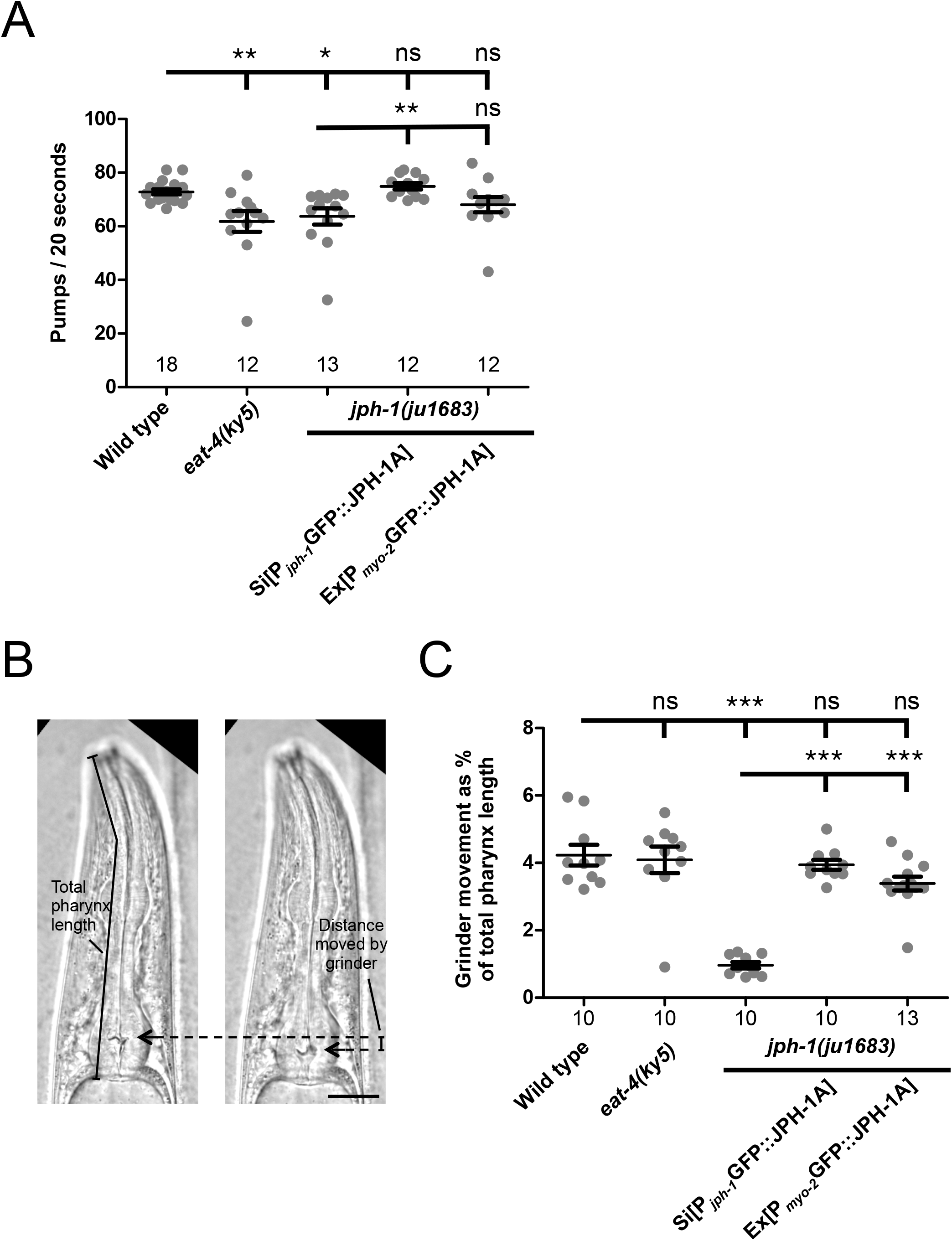
*jph-1* is required in the pharyngeal muscle for normal rate and strength of pumping. **A**) *jph-1* is required for normal pharyngeal pumping rate. *jph-1(ju1683)* mutants had reduced pumping rate, which was rescued by expression of JPH-1A by the *jph-1* promoter *[Pjph-1-GFP::JPH-1A(juSi387)]* but not by expression in the pharyngeal muscle [P*myo-2-*GFP::JPH-*1A(juEx8041)]. eat-4(ky5)* loss-of-function mutants had reduced pumping rate, as previously reported (Lee et al., 1999). Number of animals per genotype indicated above X-axis tick marks. Data are shown as individual data points and mean±SEM. Statistics: Non-parametric Kruskal-Wallis test with Dunn’s multiple comparison test. ns not significant, *p<0.05, **p<0.01. **B**) Pumping strength was determined by the distance moved by the grinder. The image on the left shows the head of the animal just before the pump is initiated, with the grinder position indicated by the arrow. The image on the right shows the animal mid-pump when the grinder has moved to its fullest extent. The distance moved by the grinder between the two images was normalized to the total length of the pharynx to quantify pumping strength. Scale bar, 25 μm. **C**) Quantification of pharyngeal pumping strength. *jph-1(ju1683)* mutants had substantially reduced grinder movement, which was rescued by expression of JPH-1A by the *jph-1* promoter *[Pjph-1-GFP::JPH-1A(juSi387)]* or in the pharyngeal muscle [P*myo-2-*GFP::JPH-1A(*juEx8041*)]. Number of animals per genotype indicated below X-axis tick marks. Data are shown as individual data points and mean±SEM. Statistics: One-way ANOVA with Tukey’s post-test. ns not significant, ***p<0.001.

### *jph-1* is required in the body wall muscle for locomotion

On solid surfaces, wild-type *C. elegans* crawl by sinusoidal body undulations (**Figure 4A**). In contrast, *jph-1(0)* mutants adopt unusual extended or curled postures during locomotion, move slowly, and are frequently immobile, consistent with previous observations of *jph-1* RNAi treated animals (Yoshida et al., 2001). When placed in liquid, *C. elegans* swim by moving their entire bodies side-to-side to produce alternating C-shaped conformations (Gjorgjieva et al., 2014), which can be quantitated by counting thrashing frequency. We observed that *jph-1(0)* mutants exhibit far fewer thrashes per minute than wild-type animals (**Figure 4B**). Furthermore, *jph-1(0)* mutants would often thrash only the heads without moving the tail. The failure of muscle contraction to propagate to the tail suggested that *jph-1* might be required for transmission of the signal for muscle contraction. A fosmid containing genomic *jph-1* fully rescued locomotion on both solid surfaces and in liquid (**Table 1, Figure 4B**). JPH-1A driven by the *jph-1* promoter rescued locomotion defects and thrashing frequency, although not as well as the fosmid transgene (**Figure 4B**). JPH-1B did not discernably improve movement on solid surfaces, supporting the importance of the transmembrane domain for JPH-1 function. Expression of JPH-1A in body wall muscle, but not pharyngeal muscle or neurons, restored fullbody thrashing in liquid and sinusoidal movement on solid surfaces (**Figure 4B**). These data indicate that *jph-1* is required in the body wall muscle for animal movement and suggest that it may be involved in both muscle contraction and propagation of a signal for contraction between muscle cells.

**Figure 4.**
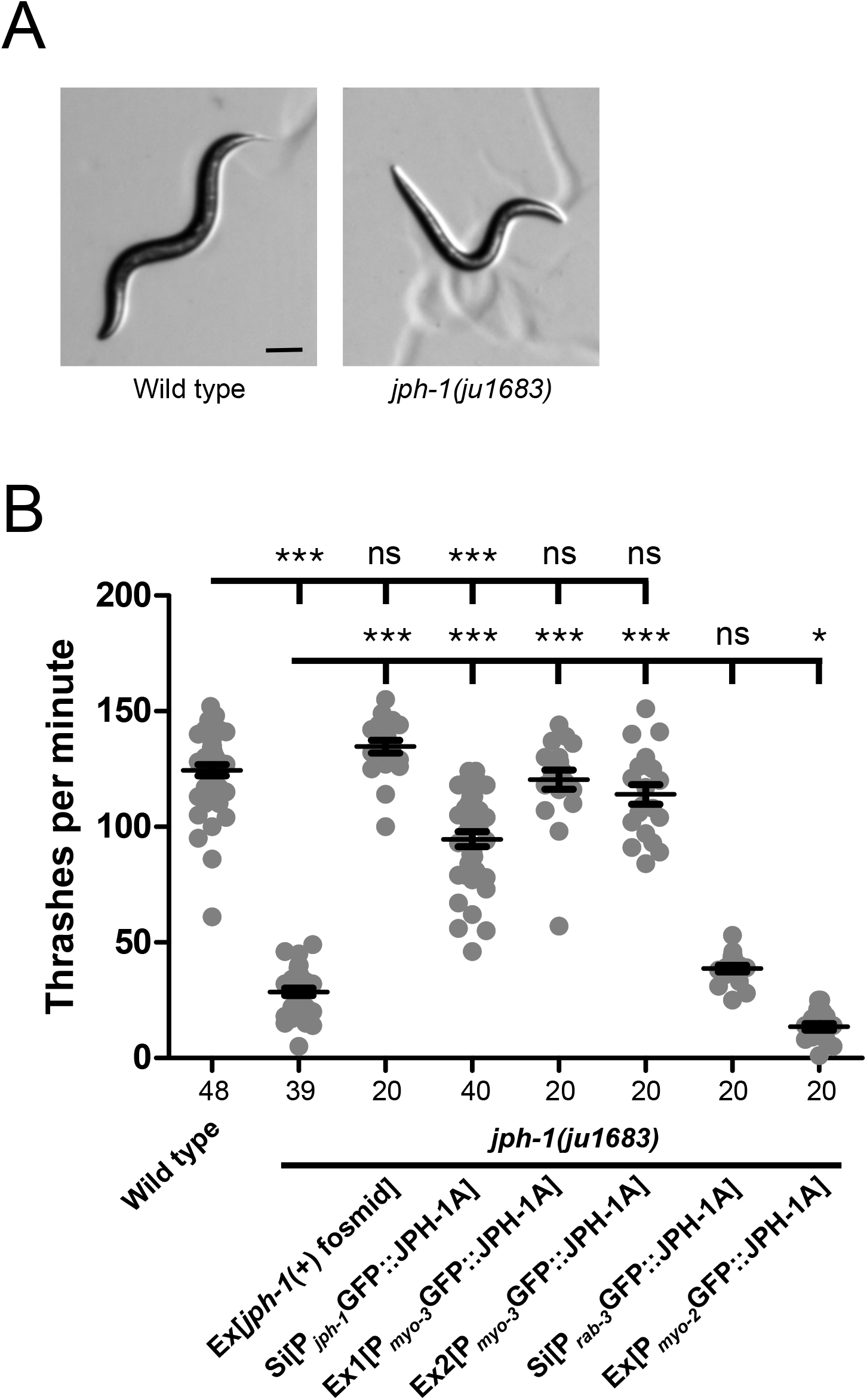
*jph-1* is required in the body wall muscle for locomotion. **A**) L4 stage wild type animals exhibit smooth sinusoidal movement and posture while *jph-1(ju1683)* animals assume unusually straight body positions (shown here) and unusually tight sinusoidal or curled positions. Scale bar, 100 μm. **B**) *jph-1(ju1683)* null mutants thrash less frequently than wild-type N2 animals. Thrashing rate was rescued by expression of a fosmid containing *jph-1 (juEx3390)* and partially rescued by expression of JPH-1A by the *jph-1* promoter *[Pjph-1-GFP::JPH-1A(juSi387)].* Expression of JPH-1A in body wall muscle [P*myo-3*-GFP::JPH-1A(*juEx8023*)] rescued thrashing rate, but expression in neurons [P*rab-3*-GFP::JPH-1A(*juSi389*)] did not. Expression of JPH-1A in the pharyngeal muscle [P*myo-2-*GFP::JPH-1A(*juEx8041*)] slightly decreased thrashing rate. Number of animals per genotype indicated below X-axis tick marks. Data are shown as individual data points and mean±SEM. Statistics: One-way ANOVA with Tukey’s post-test. ns not significant, ***p<0.001.

### *jph-1* promotes axon regeneration cell non-autonomously

We previously characterized a different *jph-1* mutation, *jph-1(ok2823),* for its role in axon regeneration. *jph-1(ok2823)* is a small deletion removing part of the fourth intron to the sixth exon (**Figure 1A**). By analyzing cDNA isolated from *jph-1(ok2823)* animals, we found that *jph-1(ok2823)* would generate a protein truncated after the seventh MORN repeat. The gross morphology and movement of *jph-1(ok2823)* animals are similar to *jph-1(0),* and these defects are fully rescued by the *jph-1* fosmid transgene (**Table 1**). We had observed that PLM axons of *jph-1(ok2823)* animals display reduced axon regeneration and enhanced axon-axon fusion after laser-induced axon injury (Kim et al., 2018). We tested if PLM axon regeneration is similarly affected in *jph-1(0)* mutants. Like *jph-1(ok2823)* animals, touch receptor neurons of *jph-1(0)* mutants have normal morphology (**Supplemental Figure 3A**), indicating that *jph-1* is not required for axon outgrowth during development. After laser injury, *jph-1(0)* mutants exhibited strongly reduced axon regeneration, significantly different from both wild type and *jph-1(ok2823)* (**Figure 5A,B**). Expression of JPH-1A under the *jph-1* promoter fully rescued the regeneration defect, indicating that *jph-1* is required for axon regrowth after injury. Expression of JPH-1A in pharyngeal muscle, which rescued the growth and size of the animal (**Table 1**), also rescued axon regrowth (**Figure 5B**), suggesting that nutrient intake may influence axon regeneration. While *jph-1* is expressed in PLM neurons (**Figure 2**), expression of JPH-1A specifically in touch neurons did not rescue axon regrowth (**Figure 5B**). Furthermore, knocking down GFP:JPH-1A specifically in touch neurons of *jph-1(0)* animals through Degron-mediated degradation of GFP-JPH-1 (Wang et al., 2017) did not reduce axon regeneration (**Supplemental Figure 3B**). Together, these data indicate that *jph-1* regulates axon regeneration cell non-autonomously.

**Figure 5.**
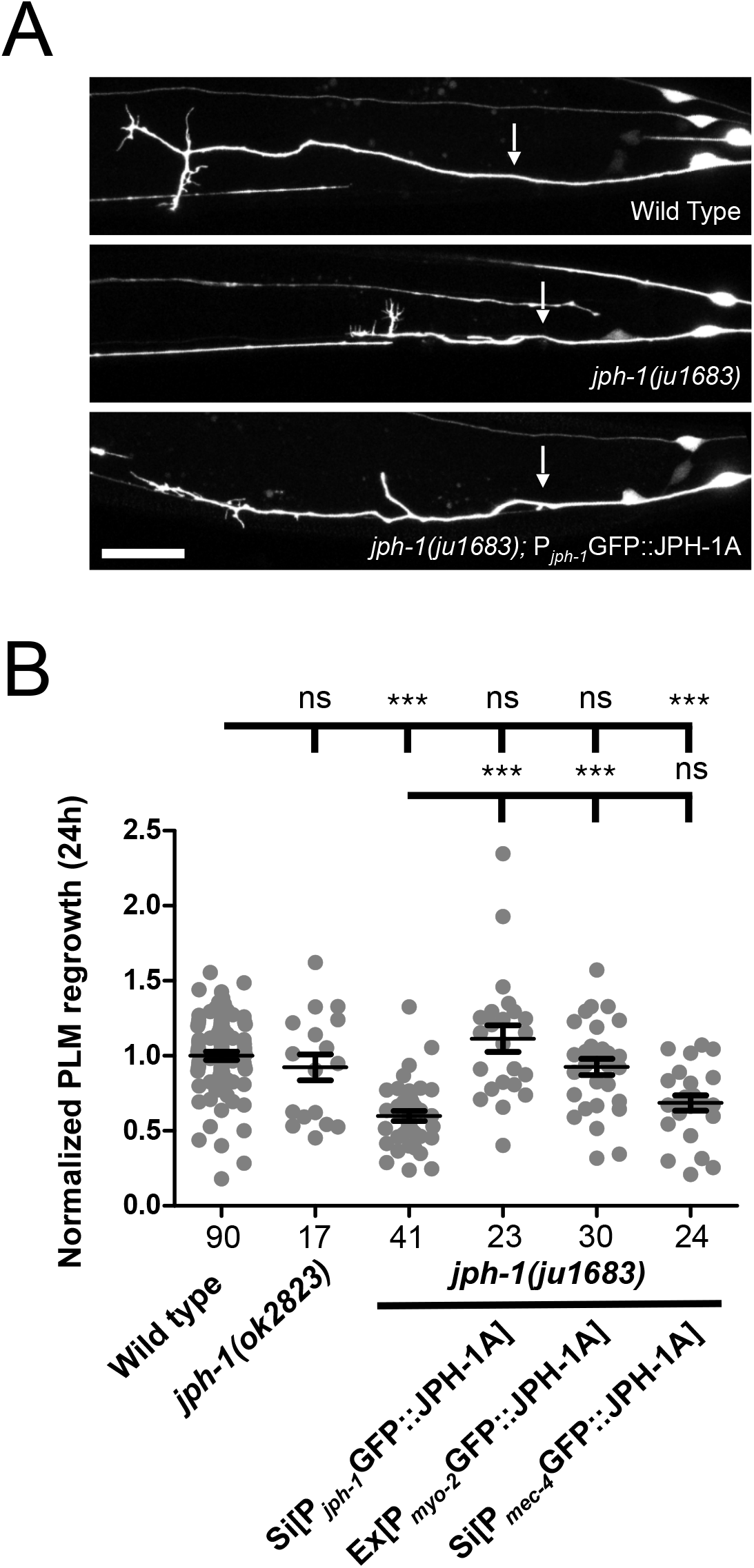
*jph-1* promotes touch neuron PLM axon regeneration cell non-autonomously. **A**) Representative confocal images of PLM axon regrowth 24 h post-axotomy in animals expressing the touch neuron marker *Pmec-7-GFP(muIs32).* Genotype in the bottom image is *jph-1(ju1683); Pjph-1-GFP::JPH-1A(juSi387).* Anterior is to the left, dorsal is up. Arrows indicate the site of axon injury. Scale bar, 20 μm. **B**) *jph-1* is required in the pharyngeal muscle for touch neuron axon regeneration. Distance regrown by PLM axon 24 h post-injury, normalized to wild-type regrowth. *jph-1(ok2823)* axon regrowth was not significantly different from wild type [P*mec-7*-GFP(*muIs32*)]. *jph-1(ju1683)* animals had significantly reduced regrowth. Expression of JPH-1A by the *jph-1* promoter *[Pjph-1-GFP::JPH-1A(juSi387)]* or in the pharyngeal muscle [P*myo-2-*GFP::JPH-1A(*juEx8041*)] rescued the reduced regrowth of *jph-1(ju1683)* mutants. Expression of JPH-1A in the touch receptor neurons [P*mec-4*-GFP::JPH-1A(*juSi388*)] did not rescue axon regeneration. Number of animals per genotype indicated below X-axis tick marks. Data are shown as individual data points and mean±SEM. Statistics: Non-parametric Kruskal-Wallis test with Dunn’s multiple comparison test. ns not significant, ***p<0.001.

While we were able to replicate the increased axon fusion of *jph-1(ok2823)* mutants, we did not observe an increase in axon fusion in injured PLM axons in *jph-1(0)* mutants (**Supplemental Figure 3C**). We considered if the enhanced axon fusion observed in *jph-1(ok2823)* animals might be caused by the production of an abnormal protein. To test this, we made a construct fusing GFP to *jph-1* cDNA isolated from *jph-1(ok2823)* animals, named GFP::JPH-1(ok2823). In contrast to the subcellular punctate pattern of full-length JPH-1A, GFP::JPH-1(ok2823) was found in the nucleus of many neurons and body wall muscles (**Supplemental Figure 3D**). Therefore, two explanations can be made for the increased axon fusion of *jph-1(ok2823)* mutants: either that *jph-1(ok2823)* is a partial loss of function and that fusion is more likely when axon regrowth is only mildly impaired, or that *jph-1(ok2823)* produces a protein with altered activity that enhances axon fusion.

### *jph-1* contributes to neuromuscular synaptic transmission

Junctophilins are required for proper regulation of cytosolic calcium levels in cell types such as mouse cardiomyocytes, HL-1 immortalized mouse cardiomyocytes, and C2C12 myotubes (Chen et al., 2013; Landstrom et al., 2011; Nakada et al., 2018; Reynolds et al., 2013; Takeshima et al., 2000; Van Oort et al., 2011). We observed broad expression of *jph-1* in neurons. Within the ventral nerve cord, we found that cholinergic motor neurons express *jph-1* (**Supplemental Figure 4A**). JPH-1A is present at the presynaptic terminal of touch receptor neurons (**Supplemental Figure 4B**). To examine if *jph-1* plays a role in synaptic transmission, we focused our study on the neuromuscular junction, where pharmacological assays can assess neuromuscular transmission. Release of acetylcholine from ventral cord motor neurons stimulates body wall muscle contraction in *C. elegans* (Von Stetina et al., 2006). Two pharmacological responses are widely used to assess neuromuscular transmission. Levamisole is an agonist of acetylcholine receptors expressed on the body wall muscle (Lewis et al., 1980). Upon exposure to 1 mM levamisole, *jph-1(0)* mutants paralyzed at the same rate as wild-type animals (**Supplemental Figure 5A**), suggesting that *jph-1* is not required for muscle responses to acetylcholine. The acetylcholinesterase inhibitor aldicarb causes the accumulation of acetylcholine at the neuromuscular junction, which leads to muscle hypercontraction and paralysis (Miller et al., 1996). Nearly all wild-type animals were paralyzed after 2 hours of exposure to 1 mM aldicarb (**Figure 6A**). In contrast, 70-80% of *jph-1(0)* mutants were still moving, suggesting that these animals may have decreased acetylcholine release. Aldicarb resistance was confirmed using a second *jph-1(0)* allele (**Supplemental Figure 5B**) and expression of a fosmid containing *jph-1* genomic DNA rescued the aldicarb resistance of *jph-1(0)* mutants (**Figure 6A**). Altogether, these results indicate that *jph-1* contributes to neuromuscular synaptic transmission.

**Figure 6.**
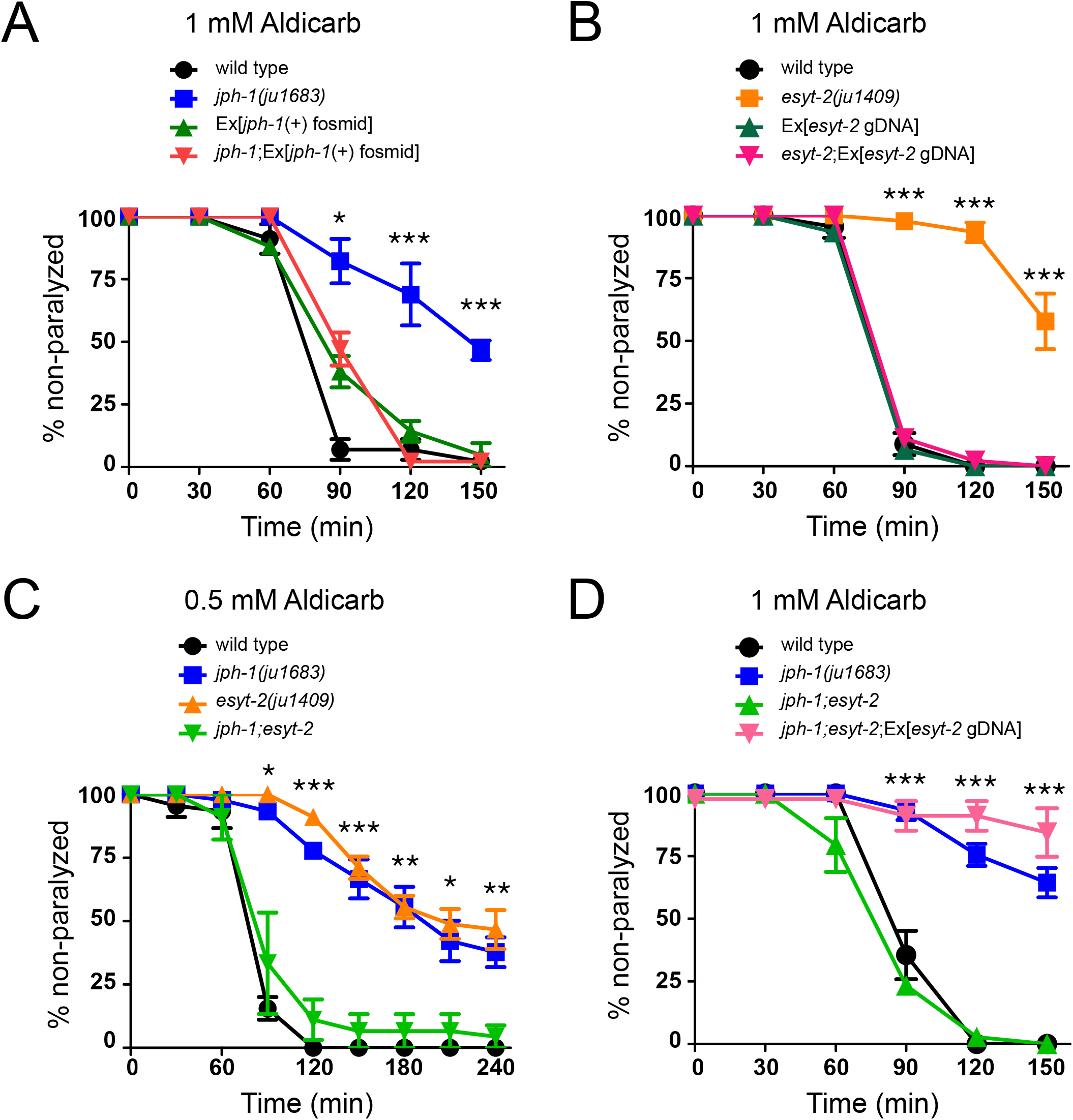
*jph-1* and *esyt-2* null mutants are aldicarb resistant and exhibit mutual suppression. **A**) *jph-1(ju1683)* animals are resistant to aldicarb compared to wild-type animals. Aldicarb resistance was rescued by expression of a fosmid containing *jph-1* genomic DNA *(juEx3390).* Statistical significance shown between *jph-1(ju1683)* and *jph-1;Ex[jph-1(+) fosmid].* **B**) *esyt-2(ju1409)* animals are resistant to aldicarb compared to wild-type animals. Aldicarb resistance was rescued by expression of *esyt-2* genomic DNA *(juEx7581).* Statistical significance shown between *esyt-2(ju1409)* and *esyt-2;Ex[esyt-2 gDNA].* **C**) *jph-1(ju1683);esyt-2(ju1409)* double mutants exhibit a wild-type response to aldicarb. Statistical significance shown between *jph-1(ju1683)* and *jph-1;esyt-2.* **D**) Expression of *esyt-2* genomic DNA *(juEx7581)* restores aldicarb resistance to *jph-1(ju1683); esyt-2(ju1409)* double mutants. Statistical significance shown between *jph-1;esyt-2* and *jph-1;esyt-2*;*Ex[esyt-2 gDNA]*. 13-15 animals tested per genotype per trial, n=3 trials. Data are shown as individual data points and mean±SEM. Statistics: One-way ANOVA with Tukey’s post-test. ns not significant, *p<0.05, **p<0.01, ***p<0.001.

As we had observed co-localization between JPH-1 and ESYT-2 in neurons, we tested the response of *esyt-2(0)* mutants to aldicarb. We found that *esyt-2(0)* mutants are aldicarb resistant, suggesting that they are also involved in neuromuscular synaptic transmission (**Figure 6B**). A transgene containing the whole *esyt-2* genomic locus rescued the aldicarb resistance of *esyt-2(0)* (**Figure 6B**). Remarkably, the *jph-1(0);esyt-2(0)* double mutant paralyzed at a similar rate to wild-type – in effect, the *jph-1(0)* and *esyt-2(0)* mutations cancel each other out (**Figure 6C**). We tested second alleles of *jph-1(0)* and *esyt-2(0)* and observed the same result (**Supplemental Figure 5C**). A transgene containing the *esyt-2* genomic locus in the *jph-1(0);esyt-2(0)* double mutant restored aldicarb resistance, indicating that the wild-type aldicarb response is due to loss of *esyt-2* (**Figure 6D**). While *esyt-2(0)* animals are superficially wild-type, *jph-1(0); esyt-2(0)* mutants resemble *jph-1(0)* in growth and locomotion, suggesting that the *esyt-2* mutation does not compensate for the loss of *jph-1* in muscles. Taken together, these results suggest that while loss of *jph-1* or *esyt-2* alone disrupts neurotransmission, loss of both restores neurotransmission to wild-type levels.

*esyt-2(0)* mutants displayed a slight resistance to levamisole that was not observed in *jph-1(0)* or *jph-1(0);esyt-2(0)* mutants (**Supplemental Figure 5D**). A non-wild type response to levamisole typically suggests a role in the muscle response to acetylcholine. However, as we had previously shown that a *esyt-2* promoter::GFP transgene is expressed exclusively in the nervous system (Kim et al., 2018), this hints that the role of *esyt-2* in neurotransmission may be more complex.

### *jph-1* promotes animal health and development in parallel with *unc-68/RyR* and voltagegated calcium channels

Our observation that JPH-1A co-localizes with UNC-68/RyR and the VGCC α1-subunit EGL-19 raises the possibility of direct interaction between them. We thus next investigated genetic interactions between *jph-1* and calcium channels in *C. elegans.*

Like *jph-1(0)* mutants, *unc-68(e540)* null mutants are small, slow growing, and show incomplete flaccid paralysis (Maryon et al., 1996). However, *unc-68(0)* mutants have darker pigmentation and grow more quickly than *jph-1(0)* mutants (**Table 2**), suggesting that they have less severe defects in nutrient intake (Avery, 1993). We found that *jph-1(0); unc-68(0)* double mutants grew even slower than either *jph-1(0)* or *unc-68(0)* single mutants (**Table 2**). Expressing JPH-1A under the *jph-1* promoter in *jph-1(0); unc-68(0)* double mutants partially restored animal growth to more closely resemble *unc-68(0)* single mutants. The exacerbated slow growth of the *jph-1(0); unc-68(0)* double mutant indicates that *jph-1* has functions independent of *unc-68* and suggests that *jph-1* may couple other ER and PM components.

**Table 2.** Summary of growth rates and movement of double mutants of *jph-1(0)* with calcium channels and *esyt-2* mutants.

| Genotype | Days to L4 (20°C) | Movement |
| --- | --- | --- |
| *jph-1(ju1683)* | 3 to 4 | Partial flaccid paralysis |
| *unc-68(e540)* | 2 to 3 | Partial flaccid paralysis |
| *egl-19(ad1006lf)* | 2 | Partial flaccid paralysis |
| *egl-19(ad695gf)* | 2 | Normal |
| *unc-2(e55)* | 2 | Paralyzed |
| *unc-2(zf35gf)* | 2 | Hyperactive |
| *esyt-2(ju1409)* | 2 | Normal |
| *jph-1(ju1683); unc-68(e540)* | 4 to 6 | Severe flaccid paralysis |
| *jph-1(ju1683); unc-68(e540);*<br>*juEx7999 [Pjph-1-GFP::JPH-1A]* | 3 to 4 | Partial flaccid paralysis |
| *jph-1(ju1683); egl-19(ad1006lf)* | Lethal |  |
| *jph-1(ju1683); egl-19(ad695gf)* | 4 to 5 | Partial flaccid paralysis |
| *jph-1(ju1683); unc-2(e55)* | 4 to 7 | Paralyzed |
| *jph-1(ju1683); unc-2(zf35gf)* | 5 to 7 | Partial flaccid paralysis |
| *jph-1(ju1683); esyt-2(ju1409)* | 3 to 5 | Partial flaccid paralysis |

All alleles are null unless otherwise annotated as gain-of-function (gf) or partial loss-of-function (lf).

*egl-19* is expressed in both muscles and neurons, and *egl-19* null mutants are embryonic lethal (Lee et al., 1997). We therefore used a partial loss-of-function mutation, *egl-19(ad1006lf),* to test genetic interactions with *jph-1.* Animals homozygous for the *egl-19(ad1006lf)* mutation are long, thin, and flaccid, move slowly, and display weak pumping (Lee et al., 1997). We were unable to obtain viable *jph-1(0); egl-19(lf)* double mutants, suggesting that *jph-1* becomes crucial when *egl-19* function is impaired. We also constructed double mutants of *jph-1(0)* with the gain-of-function mutation *egl-19(ad695gf).* Animals with *egl-19(ad695gf)* are short due to body wall muscle hypercontraction (Lainé et al., 2014) but otherwise appear normal in overall growth rate and movement. We found that *jph-1(0); egl-19(gf)* animals lived to adulthood, but grew more slowly than *jph-1(0)* single mutants (**Table 2**). Overall, these observations suggest that when *egl-19* activity is impaired or altered, *jph-1* activity becomes more important.

The non-L-type VGCC α1-subunit *unc-2,* orthologous to CACNA1A, is predominantly expressed in neurons and localizes to presynaptic terminals (Mathews et al., 2003; Saheki and Bargmann, 2009)*. unc-2(e55)* null mutants exhibit sluggish movement but normal development and growth (Mathews et al., 2003; Schafer et al., 1996). We found that *jph-1(0); unc-2(0)* double mutants grew substantially more slowly than *jph-1(0)* single mutants and were sterile as adults (**Table 2**). The *unc-2(zf35gf)* gain-of-function mutation causes the channel to open at a lower membrane potential, causing hyperactive locomotion but otherwise normal growth and development (Huang et al., 2019). *jph-1(0); unc-2(gf)* double mutants displayed significantly slower growth than *jph-1* single mutants (**Table 2**). These results suggest that *jph-1* and *unc-2* function cooperatively in neurons.

Altogether, our analysis of genetic interactions supports a conclusion that *jph-1* acts together with RyR and VGCC channels for animal development, where they are not in completely overlapping pathways but may have some overlapping roles.

### JPH-1A subcellular localization depends on *unc-68/RyR*

Evidence from other cell types suggest that junctophilins and their interacting partners may depend on each other to be localized to MCS (Golini et al., 2011; Nakada et al., 2018). We thus tested if JPH-1A localization depends on calcium channels and *esyt-2.* In the body wall muscle of wild type animals, JPH-1A localizes to longitudinal rows of puncta (**Figure 7A**). In *unc-68* mutants, JPH-1A puncta were less distinct and often connected to neighbouring puncta. (**Figure 7A**). In wild-type neurons, JPH-1A has a reticulate pattern with bright puncta in the cell periphery (**Figure 7B**). In *unc-68* mutants, while the reticulate pattern of JPH-1 remained, the bright puncta were absent (**Figure 7B**). The lack of puncta in both muscles and neurons of *unc-68* animals suggests that *unc-68* is required for anchoring JPH-1A in puncta. JPH-1A localization was unchanged from wild type in *unc-2* and *esyt-2* mutants (**Figure 7, Supplemental Figure 6A-B**), indicating that these genes are not required for JPH-1A localization.

**Figure 7.**
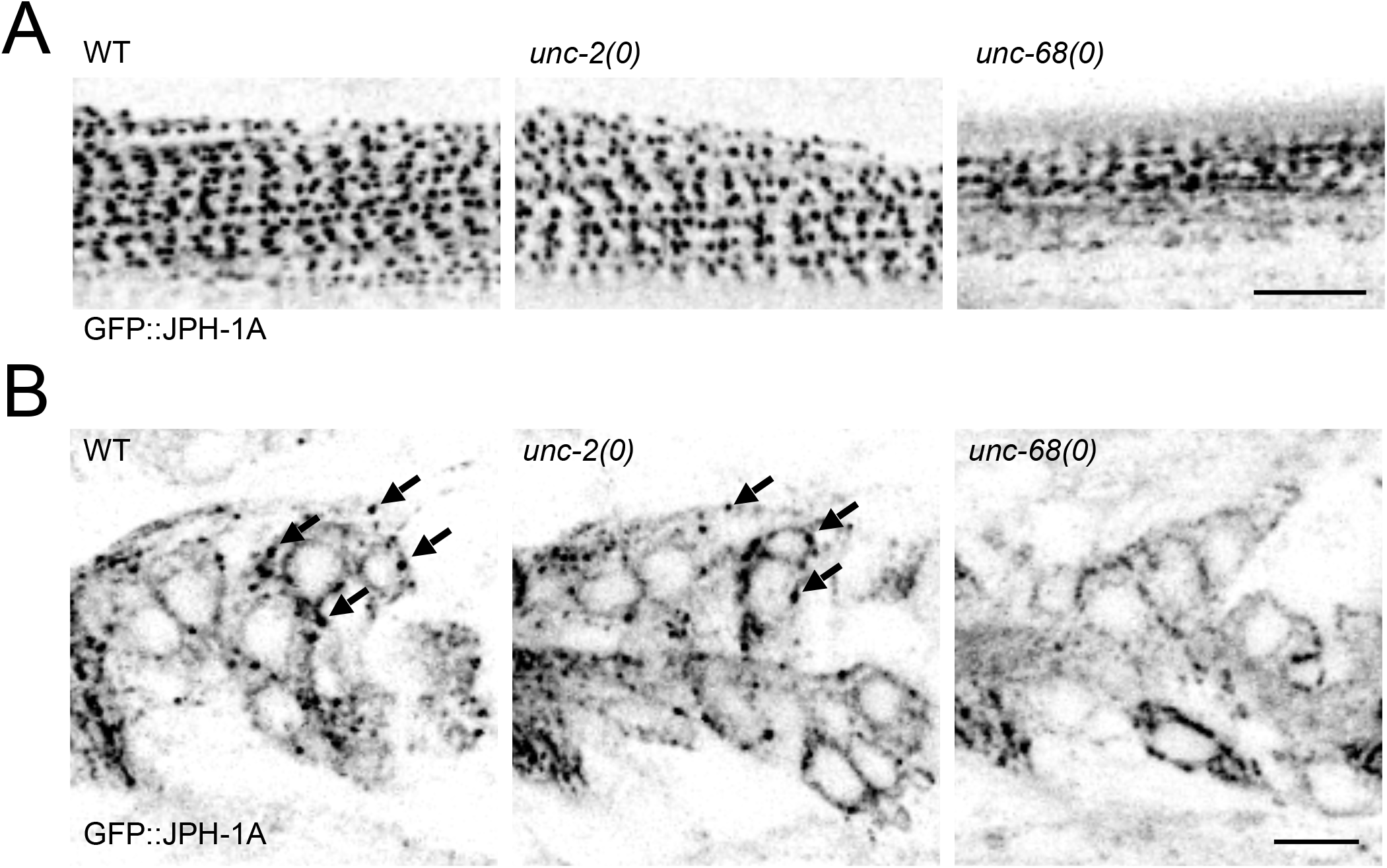
*unc-68* is required for JPH-1A localization. Shown are single-plane confocal images of GFP::JPH-1A expressed under the *jph-1* promoter *[Pjph-1-GFP::JPH-1A(juEx7999)]* in wild-type (WT), *unc-2(e55),* and *unc-68(e540)* backgrounds. **A**) In the body wall muscle, JPH-1A localizes to row of puncta in WT and *unc-2(e55)* animals, while in *unc-68(e540)* animals JPH-1A puncta are less distinct. **B**) In neurons of the head ganglia, JPH-1A localizes to reticulate structures surrounding the nucleus and forms puncta in the cell periphery of WT and *unc-2(e55)* animals, but not *unc-68(e540)* mutants. Arrows mark some of the GFP::JPH-1A puncta. WT and *unc-2(e55)* images were taken at 2% laser power and *unc-68(e55)* was taken at 4.5% laser power to compensate for the slight variation in expression level. Scale bar, 5 μm.

## Discussion

Junctophilins play key roles in excitation-contraction coupling in heart and skeletal muscles (Ito et al., 2001; Nakada et al., 2018; Takeshima et al., 2000; Van Oort et al., 2011). In particular, junctophilins couple PM- and ER-localized calcium channels to efficiently trigger calcium release from the ER following membrane depolarization (Chen et al., 2013; Nakada et al., 2018; Reynolds et al., 2013; Van Oort et al., 2011). Here, we report that the *C. elegans* junctophilin JPH-1 is expressed in pharyngeal muscle, body wall muscle, and neurons, and performs important functions in each tissue. We show that in the pharyngeal muscle, *jph-1* is required for robust pumping and timely growth and development. The stunted development of *jph-1(0)* mutants is likely due to reduced food intake caused by weak pumping, as their slow growth and starved appearance is seen in other mutants with defects in feeding related function (Avery, 1993; Avery and Horvitz, 1989). In the body wall muscle, we find that *jph-1* is required for body movement and locomotion. *jph-1(0)* mutants move slowly and display flaccid paralysis, suggesting that the body wall muscle lacks contraction strength. Our tissue-specific rescue experiments indicate that muscle contraction in both pharyngeal and body wall muscle requires *jph-1.* In flies, knockdown or overexpression of the sole junctophilin was shown to cause muscular deficits and cardiac dysfunction (Calpena et al., 2018). Skeletal muscle from neonatal JPH-1 knockout mice have weaker electrically-stimulated contractile force, indicating that JPH-1 is required for excitation-contraction coupling (Ito et al., 2001). Thus, the role for junctophilin in muscle contraction is conserved from *C. elegans* pharyngeal and body wall muscle to vertebrates.

The role of calcium regulation in axon regeneration in *C. elegans* has been widely demonstrated (Ghosh-Roy et al., 2010). *unc-68/RyR* promotes axon regeneration, and is required for localized calcium release from the ER following axon injury (Sun et al., 2014). We previously reported that *jph-1(ok2823)* mutants have decreased axon regeneration (Kim et al., 2018). Here, we extended our analysis to the genetic null alleles of *jph-1* and uncovered a surprising role of *jph-1* in promoting axon regeneration in a cell non-autonomous manner. The observation that the regeneration defects could be rescued by expressing *jph-1* in the pharyngeal muscle implies that PLM axon regeneration may be influenced by nutrient uptake or through substances released by the pharynx. This finding raises an intriguing possibility that gut nutrients may impact neuronal injury response, a theme that shares similarities to emerging findings on the gut-brain axis in other axon regeneration studies (Kigerl et al., 2020). Additionally, despite *jph-1(ok2823)* animals resembling *jph-1(0)* in all gross phenotypes, our data suggest that the increased fusion in *jph-1(ok2823)* is likely due to an altered activity associated with the truncated protein JPH-1(ok2823) that localizes to the nucleus. Interestingly, a study in mouse found that heart stress induces cleavage of JPH-2, with the N-terminal JPH-2 fragment translocating to the nucleus where it alters transcription (Guo et al., 2018). Therefore, it is conceivable that the mutant protein produced in *jph-1(ok2823)* alters neuronal transcription to enhance axon fusion after injury.

Our finding that *jph-1(0)* mutants are resistant to the acetylcholinesterase inhibitor aldicarb suggests that *jph-1* modulates neurotransmission at the neuromuscular junction. The fact that *jph-1(0)* mutants showed a normal response to the acetylcholine receptor agonist levamisole suggests that *jph-1* modulates neurotransmission by functioning in neurons. In JPH-3/4 double knockout mice, paired-pulse stimulation of climbing fibres elicits normal depression in Purkinje cells, but paired-pulse stimulation of parallel fibres elicits reduced facilitation in Purkinje cells, leading the authors of the study to conclude that JPH-3/4 may play a subtle role in mammalian synaptic transmission (Kakizawa et al., 2007). Our work suggests that *jph-1* may have a role in synaptic transmission that has largely been overlooked in studies on neuronal junctophilins in mammals. In hippocampal neurons, junctophilins couple PM-localized CaV1.3 VGCCs, ER-localized RyR2 Ca^2+^-gated Ca^2+^ channels, and PM-localized KCa3.1 Ca^2+^-activated K^+^ channels (Sahu et al., 2019). This coupling generates the slow afterhyperpolarization current, which regulates action potential frequency. Unlike mammalian neurons, which generate voltage-gated Na^+^ channel-dependent action potentials, *C. elegans* neurons mostly rely on a calcium current for membrane depolarization (Goodman et al., 1998). Therefore, while junctophilins likely regulate calcium-induced calcium release in both *C. elegans* and mammalian neurons, the physiological consequences of losing junctophilin depend on neuronal properties.

Our data further uncovers intriguing genetic interactions between *jph-1* and *esyt-2* in synaptic transmission. Extended-synaptotagmin was shown to have a presynaptic role in neurotransmission in *Drosophila* (Kikuma et al., 2017). Consistently, we found that *esyt-2* null mutants were aldicarb resistant. Strikingly, we found that *jph-1(0)*; *esyt-2(0)* double mutants had a wild-type response to aldicarb. This mutual suppression suggests that when either *jph-1* or *esyt-2* is mutated, neurotransmission is unbalanced; when the other is also mutated, the balance is restored. As we do not yet know whether *jph-1* and *esyt-2* function pre-or postsynaptically, the mechanism is unclear. However, as both proteins are ER-PM tethers, the mechanism likely involves ER calcium release. It would be of future interest to determine the exact nature of how *jph-1* regulates neurotransmission.

Finally, our genetic double mutant analysis sheds light on the importance of JPH-1 mediated ER-PM calcium channel coupling. Many studies on junctophilins have focused on their roles in coupling the ER-localized RyR with PM-localized channels in muscles and neurons. In *C. elegans,* RyR is encoded by *unc-68.* Early studies showed using both *unc-68* promoter::GFP and anti-UNC-68 immunostaining that *unc-68* is expressed in muscles and neurons, but absent in the anterior pharynx (Maryon et al., 1998). *unc-68* null mutants are aldicarb resistant, and electrophysiological studies have shown that *unc-68* has a pre-synaptic role in synaptic transmission (Liu et al., 2005; Maryon et al., 1998). We observed *jph-1* expression in the entire pharynx. Close comparison of *jph-1* and *unc-68* null mutants showed that they have similar movement and growth phenotypes, but *jph-1(0)* exhibited more severe growth retardation. Moreover, *jph-1(0); unc-68(0)* double mutants exhibit more severe growth defects than *unc-68* or *jph-1* single mutants. This analysis suggests that JPH-1 has additional RyR-independent roles. Possibilities include the generation of ER-PM contact sites, regulation of store-operated calcium entry (Hirata et al., 2006; Li et al., 2010), and coupling other ER components to the PM. Previous studies have shown that junctophilins are required for the co-localization of ER-and PM-localized calcium channels in isolated mouse cardiomyocytes, mouse skeletal muscle, and cultured hippocampal neurons (Nakada et al., 2018; Sahu et al., 2019; Van Oort et al., 2011). In rat cardiomyocytes, RyR localizes to muscle triads before JPH-2 arrives (Ziman et al., 2010), suggesting that the targeting of junctophilins by RyR may be conserved. Junctophilins and RyRs have been shown to directly interact (Beavers et al., 2013; Golini et al., 2011; Nakada et al., 2018; Phimister et al., 2007; Van Oort et al., 2011; Woo et al., 2008). We found that JPH-1 localization depends on *unc-68*/RyR. It is possible that junctophilin targeting may involve directly binding to RyR already localized at MCSs.

In conclusion, our study shows that *C. elegans jph-1,* similar to vertebrate homologs, has broad functions in excitable cells. Our data uncover new roles of junctophilins in synaptic transmission and axon regeneration, and the requirement for RyR in junctophilin localization. The conservation in function between mammalian and *C. elegans* junctophilins presents the opportunity for *C. elegans* to be used for further investigations of junctophilins.

## Acknowledgements

We thank our lab members for discussion and comments. This work was supported by grants from NIH (R01 NS093588 to ADC and YJ, R01 GM054657 (ADC), R01 NS32196 to JMK, and R37 NS 035546 to YJ).

---

**Supplemental Figure 1.**
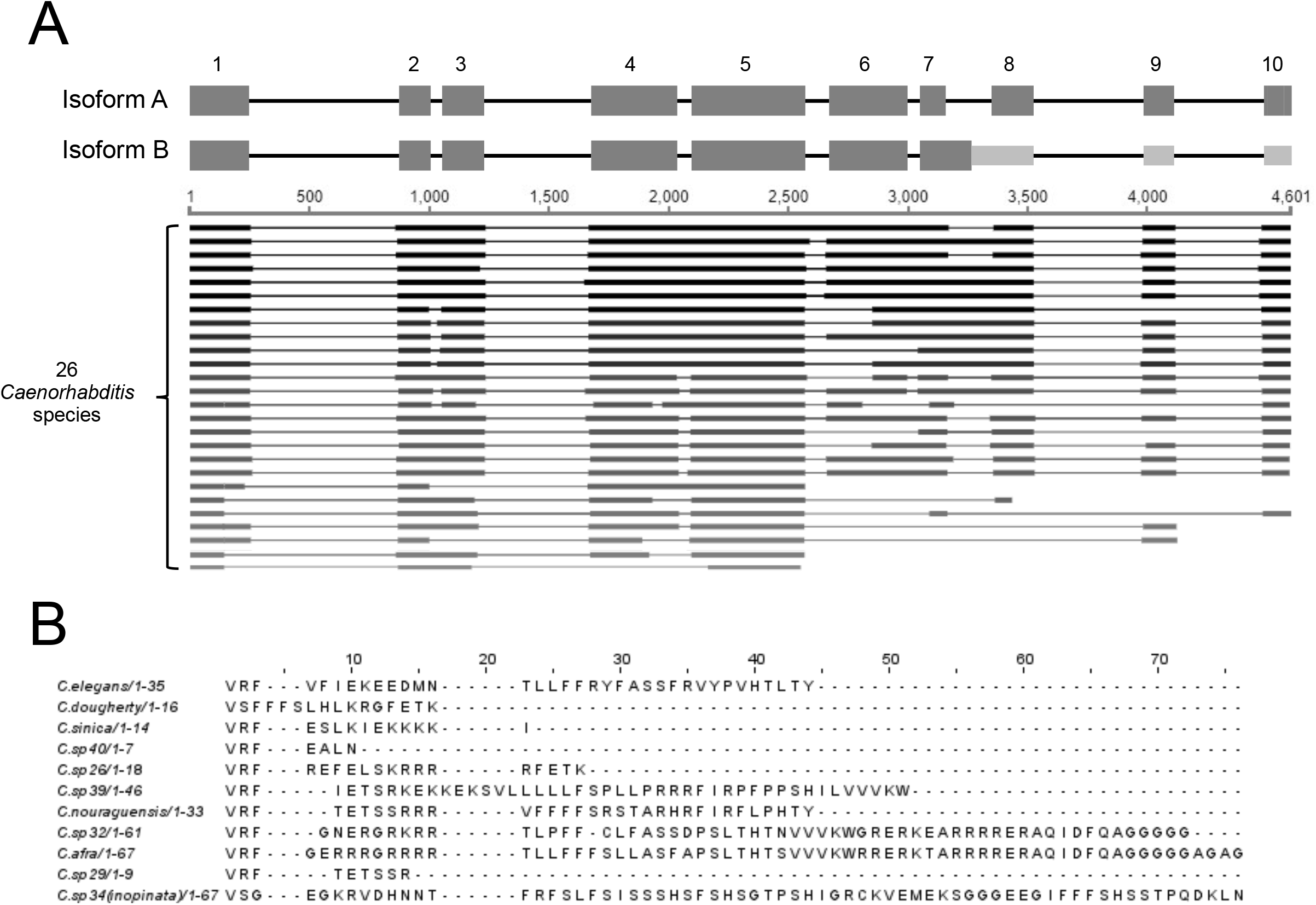
Comparison of genomic sequences concerning the intron retained in JPH-1B. **A**) Top: Exon-intron diagrams for *C. elegans jph-1* isoform A and B. Bottom: We performed a BLASTn search of *C. elegans jph-1* against 26 *Caenorhabditis* genomes published on Caenorhabditis.org. Aligned sequences are thick black lines and unaligned sequences are thin black lines. Darker lines indicate stronger hits. Boundaries between aligned and unaligned regions often match up with exon-intron boundaries. 10 *Caenorhabditis* sequences align with the intron retained in JPH-1B. **B**) We translated the introns of these 10 species in the same reading frame as *C. elegans jph-1* and aligned the amino acid sequences using MUSCLE (Madeira et al., 2019). The sequences vary in length because most encounter stop codons, except for sister species *C. sp32* and *C. afra,* which have no stop codons in the intron and are in frame with the following exon. Beyond the first three amino acids there is little amino acid conservation between sequences.

**Supplemental Figure 2.**
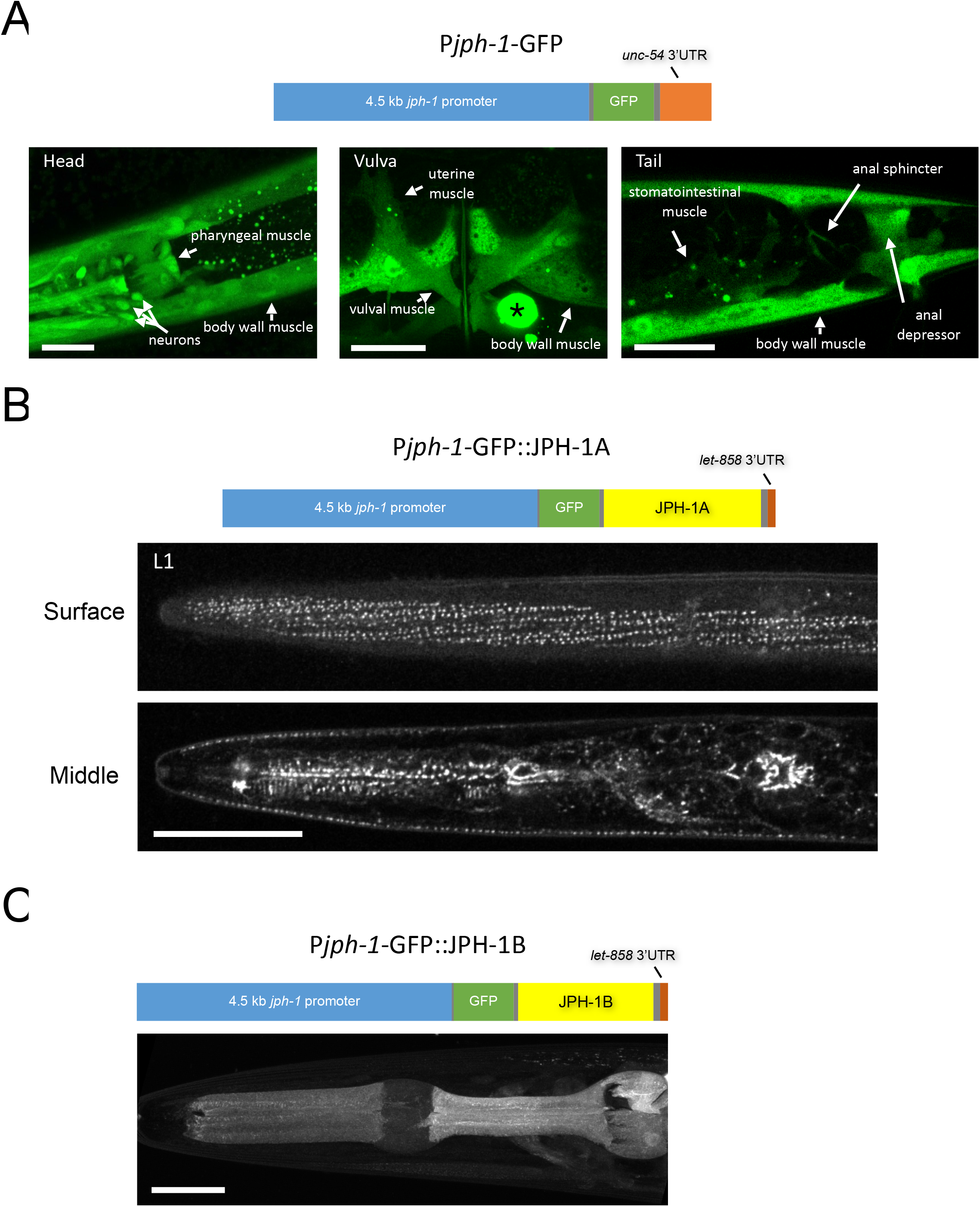
Expression pattern of a *jph-1* transcriptional reporter and a JPH-1B translational fusion reporter. **A**) *jph-1* is expressed in neurons and most muscles. Top: Illustration of *jph-1* promoter::GFP expression construct [P*jph-1*-GFP(*juEx8013* and *juEx8014*)]. Bottom: GFP expression was seen in head ganglia neurons and pharyngeal, body wall, vulval, uterine, stomatointestinal, anal sphincter, and anal depressor muscles. The large fluorescent circle marked by an asterisk is a coelomocyte labeled by the coinjection marker [P*unc-122*-RFP]. **B**) *jph-1* localization in an L1 stage animal. Top: Illustration of expression construct *[Pjph-1-GFP::JPH-1A(juSi387)].* Bottom: Confocal images of an L1 stage animal. A plane near the surface of the animal shows expression in the body wall muscle, while a plane taken through the middle of the animal shows expression in the pharyngeal muscle and head ganglia neurons. **C**) JPH-1B has a diffuse localization. Top: Illustration of construct expressing *jph-1b* cDNA under the *jph-1* promoter *[Pjph-1-GPF::JPH-1B(juEx8038)].* Bottom: Confocal projection of an L4 stage animal head shows a diffuse localization in neurons and muscles. Scale bars 20 μm.

**Supplemental Figure 3.**
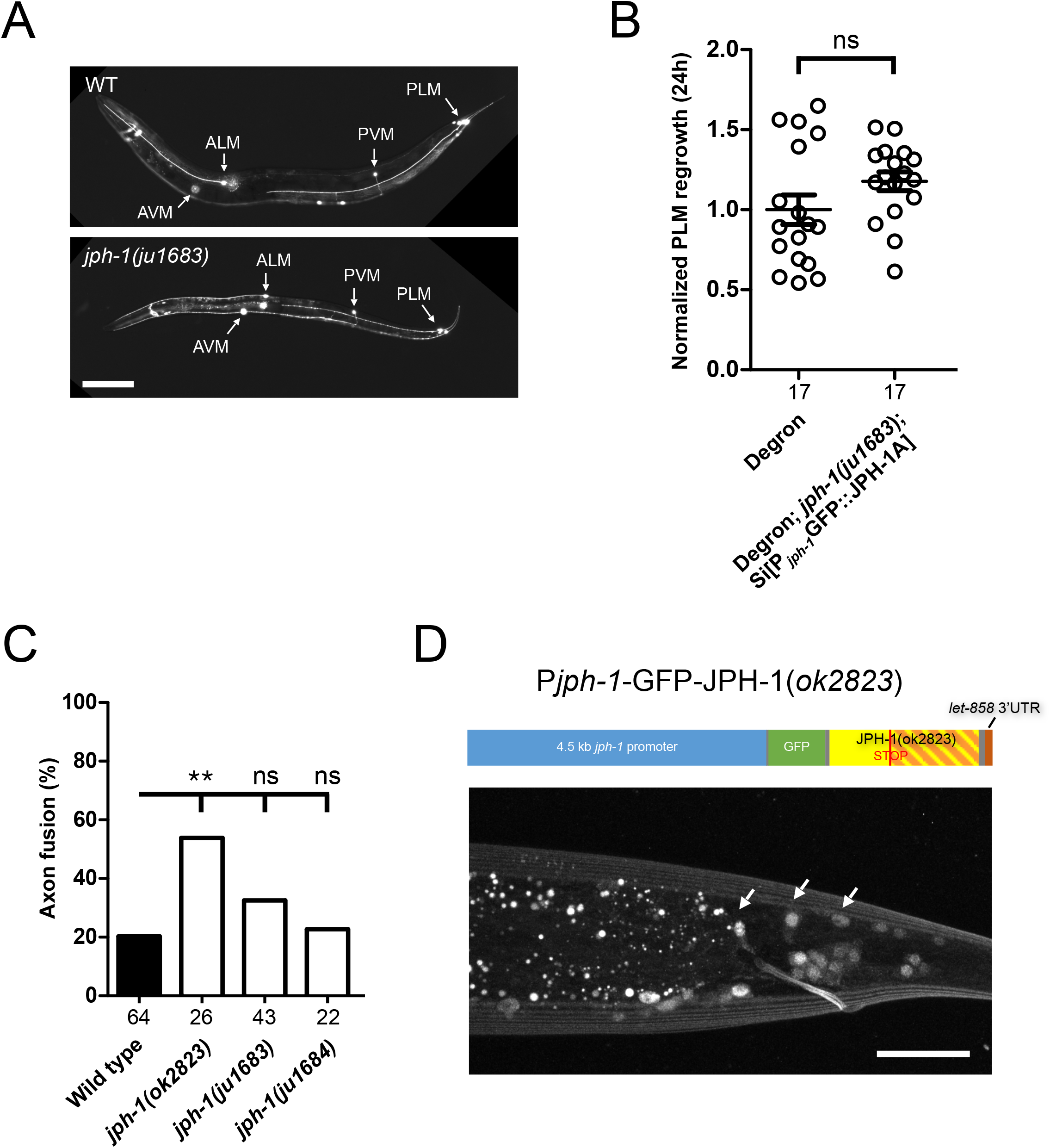
*jph-1(0)* mutants do not alter touch neuron morphology or enhance axon fusion after injury. **A**) Touch neuron morphology is normal in *jph-1(ju1683)* animals. Representative images of wildtype and *jph-1(ju1683)* day-1 adult animals expressing the touch neuron marker P*mec-7*-GFP(*muls32*). Labels indicate ALM, PLM, AVM, and PVM neuron cell bodies. The bright spot below the *jph-1(ju1683)* ALM cell body is likely fluorescence from the ALM on the opposite side of the body. Scale bar, 100 μm. **B**) Distance regrown by PLM axon 24h post-injury. Control animals expressed GFP Degron in the touch neurons. *jph-1(ju1683)* animals expressing GFP-tagged JPH-1A under the *jph-1* promoter *[Pjph-1-GFP::JPH-1A(juSi387)]* also expressed GFP Degron in the touch neurons, predicted to degrade GFP::JPH-1 specifically in touch neurons. There was no statistically significant difference between groups. Number of animals per genotype indicated below X-axis tick marks. Data are shown as individual data points and mean±SEM. Statistics: Student’s t-test. ns not significant. **C**) Percentage of animals with axon-axon fusion 24h post-injury. *jph-1(ok2823)* mutants had increased axon fusion while null mutants *ju1683* and *ju1684* exhibited wild-type levels of axon fusion. Number of animals per genotype indicated below X-axis tick marks. Statistics: Fisher’s exact test performed pairwise. ns not significant, **p<0.01. **D**) JPH-1(*ok2823*) localizes to the nucleus. Top: Illustration of construct expressing *jph-1(ok2823)* cDNA from original start to stop codon under the *jph-1* promoter [P*jph-1*-GFP::JPH-*1(ok2823)(juEx8035)].* A premature stop codon in the middle of JPH-1(*ok2823*) truncates the C-terminal two-thirds of the protein. Bottom: Confocal projection of L4-stage animal tail with arrows indicating neuronal nuclei labeled by JPH-1(ok2823). Scale bar, 20 μm.

**Supplemental Figure 4.**
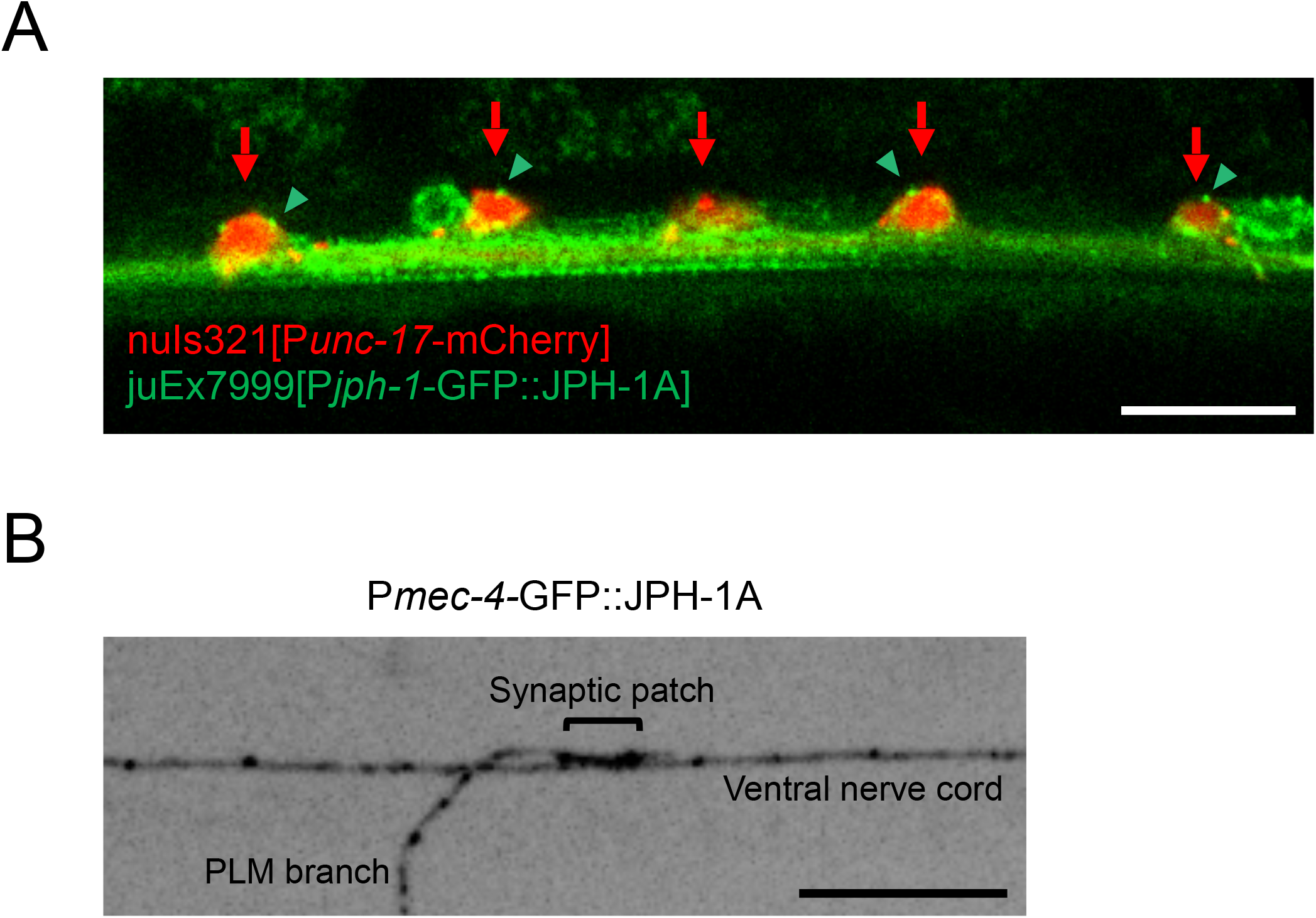
*jph-1* is expressed in cholinergic motor neurons and touch receptor neurons. **A**) *jph-1* is expressed in cholinergic neurons. Single plane confocal image of ventral nerve cord of L4 animal expressing mCherry in cholinergic neurons [P*unc-17-*mCherry(*nuIs321*)] and JPH-1A under the *jph-1* promoter *[Pjph-1-GFP::JPH-1A(juEx7999)].* Red arrows indicate cholinergic neuron cell bodies. Green arrowheads indicate JPH-1A puncta in cholinergic neurons. Scale bar, 10 μm. **B**) JPH-1A is present where the PLM touch receptor neuron synapses onto the ventral nerve cord. Confocal projection of JPH-1A expressed in touch neurons [P*mec-4*-GFP::JPH-1A(*juSi388*)]. Scale bar, 10 μm.

**Supplemental Figure 5.**
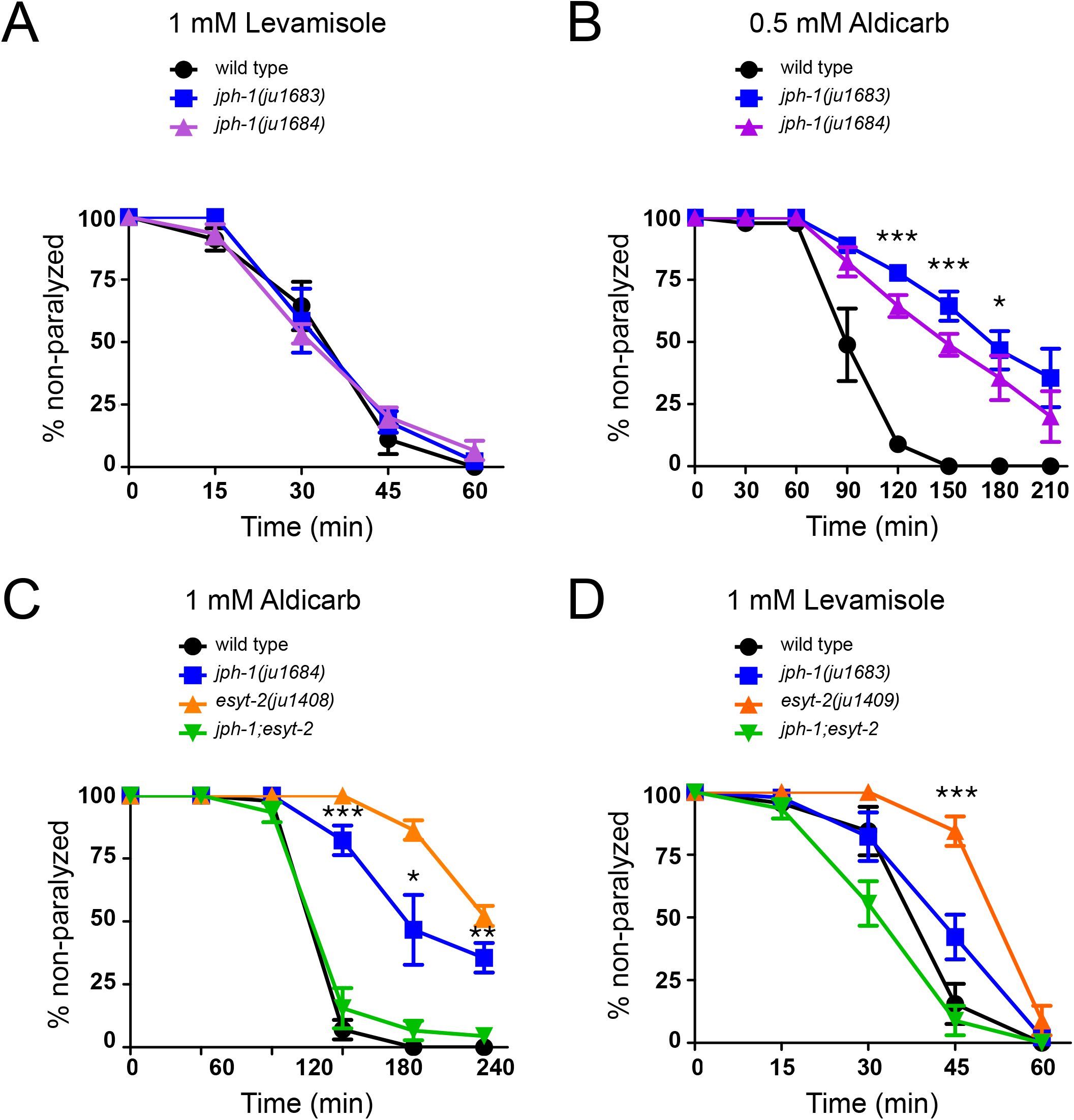
Additional data on pharmacological responses of *jph-1(0)* and *jph-1(0); esyt-2(0)* **A**) *jph-1* null mutants *ju1683* and *ju1684* have the same response to levamisole as wild type animals. **B**) *jph-1* null mutants *ju1683* and *ju1684* are both aldicarb resistant. Statistical significance shown between *jph1-(ju1684)* and wild type. **C**) *jph-1(ju1684);esyt-2(ju1408)* double mutants exhibit a wild-type response to aldicarb. Statistical significance shown between *jph-1(ju1684)* and *jph-1;esyt-2*. **D**) *esyt-2(ju1409)* animals are levamisole resistant compared to wild-type animals. Statistical significance shown between *esyt-2(ju1409)* and wild type. 13-15 animals tested per genotype per trial, n=3 trials. Data are shown as individual data points and mean±SEM. Statistics: One-way ANOVA with Tukey’s post-test. ns not significant, *p<0.05, **p<0.01, ***p<0.001.

**Supplemental Figure 6.**
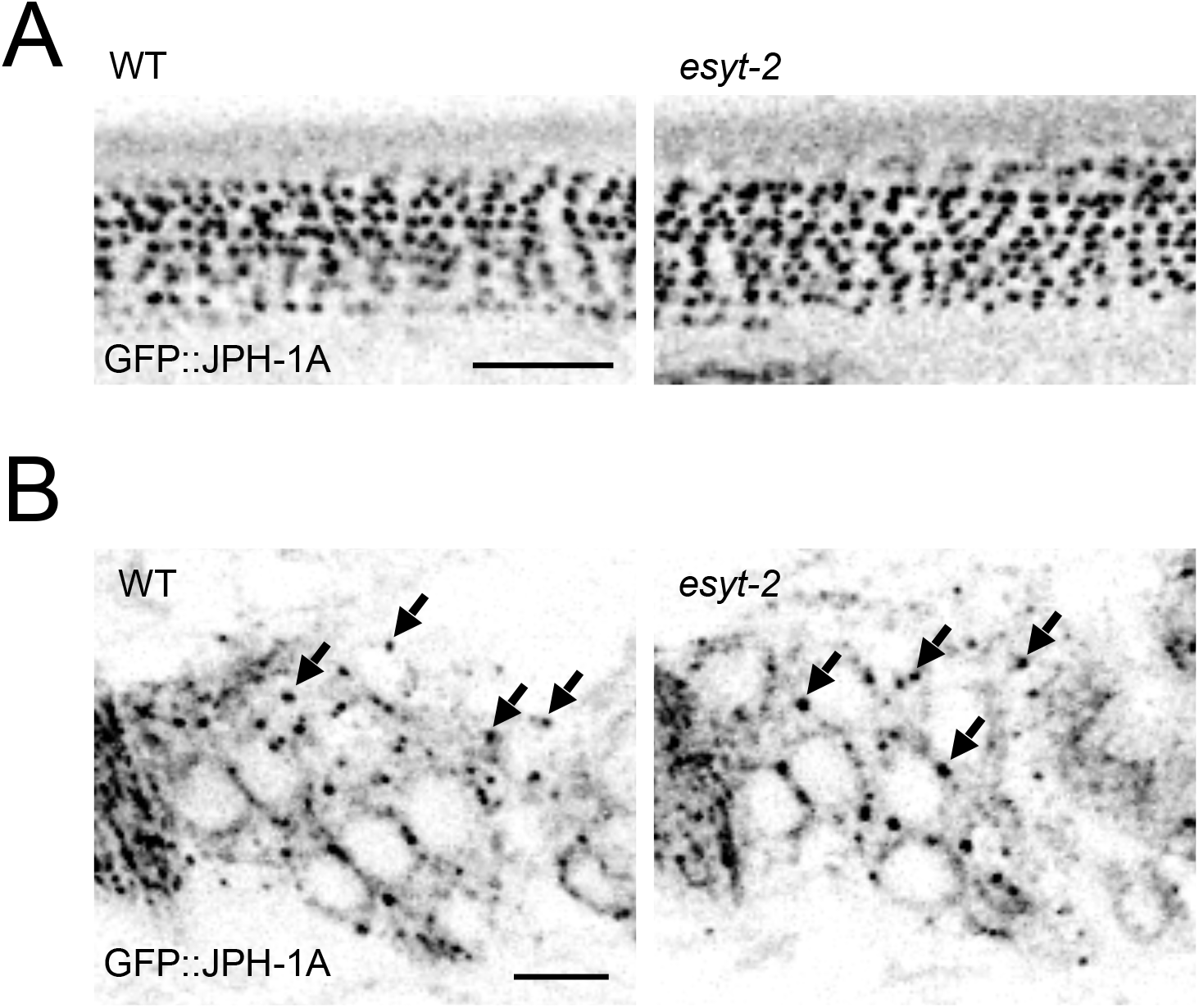
JPH-1A localization is unaltered in *esyt-2(0)* Shown are single-plane confocal images of GFP::JPH-1A expressed under the *jph-1* promoter *[Pjph-1-GFP::JPH-1A(juSi387)]* in wild-type and *esyt-2(ju1409)* backgrounds. **A**) In the body wall muscle, JPH-1A localizes to rows of puncta in wild type and *esyt-2(ju1409)* animals. **B**) In neurons of the head ganglia, JPH-1A localizes to reticulate structures surrounding the nucleus and forms puncta in the cell periphery of wild type and *esyt-2(ju1409)* animals. Arrows mark some of the GFP::JPH-1A puncta. Scale bar, 5 μm.

